# Context-invariant socioemotional encoding by prefrontal ensembles

**DOI:** 10.1101/2023.10.19.563015

**Authors:** Nicholas A. Frost, Kevin C. Donohue, Vikaas Sohal

## Abstract

The prefrontal cortex plays a key role in social interactions, anxiety-related avoidance, and flexible context-dependent behaviors, raising the question: how do prefrontal neurons represent socioemotional information across different environments? Are contextual and socioemotional representations segregated or intermixed, and does this cause socioemotional encoding to remap or generalize across environments? To address this, we imaged neuronal activity in the medial prefrontal cortex of mice engaged in social interactions or anxiety-related avoidance within different environments. Neuronal ensembles representing context and social interaction overlapped more than expected while remaining orthogonal. Anxiety-related representations similarly generalized across environments while remaining orthogonal to contextual information. This shows how prefrontal cortex multiplexes parallel information streams using the same neurons, rather than distinct subcircuits, achieving context-invariant encoding despite context-specific reorganization of population-level activity.

## Main

The medial prefrontal cortex (mPFC) plays critical roles in regulating social interactions (*1–3*) and anxiety-related behaviors (*4–7*). Both types of behaviors can occur in a variety of different locations. Some socioemotional behaviors generalize across many environments and situations, while others require specific adaptations. Information pertinent to social interactions or anxiety-related behaviors must therefore be represented in a reliable manner which is also flexible across different situational contexts.

Social interactions are encoded by single prefrontal neurons (*8, 9*), neuronal ensembles (*10, 11*), and changes in neuronal interactions(*12–15*). Underscoring the critical role of the mPFC for social interaction in mice (*16*) and humans (*17*), disrupting prefrontal computations or ensemble activity alters social behaviors (*2, 11, 16*). Similarly, the mPFC encodes information pertinent to anxiety-related states (*4, 5*), presumably reflecting the influence of anxiety-related input from sources including the ventral hippocampus (*6, 7, 18, 19*).

Both social interactions (*8*) and anxiety-related behaviors (*20*) trigger specific patterns of mPFC output. However, the mPFC also encodes other types of information including location (*8, 21, 22*) and context (*23*), which may influence how social interactions or anxiety-related behaviors are encoded when they occur at different locations or in different environments. Underscoring the challenge of understanding how contextual and socioemotional encoding interact in prefrontal cortex, recordings in the mPFC of rodents and PFC of marmosets have demonstrated that activity underlying both exploration (*23*) and social signals (specifically vocalizations) (*24*) remaps dynamically in response to changes in context, and that mouse mPFC neurons may only be active when social investigation occurs at specific locations (*8*).

We therefore sought to understand how the population level or ensemble encoding of social and anxiety-related behaviors interacts with encoding of context in the mPFC of mice exposed to distinct environments. In general, we anticipate four general possibilities: (1) socioemotional and contextual information are encoded by largely non-overlapping neuronal ensembles; (2) the same neurons encode socioemotional and contextual information but socioemotional representations are context-invariant; (3) the same neurons encode socioemotional and contextual information, but socioemotional encoding fails to generalize across contexts; (4) socioemotional encoding undergoes remapping or interacts nonlinearly with contextual information, such that different ensembles encode socioemotional information in different contexts.

To address this question, we utilized microendoscopic calcium recordings of prefrontal neurons as mice engaged in social interactions or anxiety-related behaviors across different environments. Consistent with previous reports we find that overall population-level activity within the mPFC is dynamically reorganized in a context-dependent manner. Nevertheless, the specific neuronal ensembles which strongly encode socioemotional information are context-invariant. Furthermore, representations of both social and anxiety-related information maintain a near-orthogonal orientation relative to representations of contextual information. This enables socioemotional encoding to generalize across contexts, even though the underlying socioemotional and context-encoding ensembles utilize overlapping populations of neurons.

## Results

We performed optical GCaMP measurements of prefrontal circuit activity (**Figure 1A-C**) in 8 mice. Each mouse was tested during interactions interacted with novel sex-matched juvenile conspecifics on two days. On the first day (‘static context’), all four interactions occurred within a single environment – the mouse’s home cage, and novel juveniles were introduced sequentially, interleaved with periods during which the mouse was alone (**Figure 1D**). On a separate day (‘dynamic context’), mice were first sequentially exposed to two novel juveniles in their home cage. Then they were then moved to a second context (which was familiar to them) where two additional novel juvenile conspecifics were sequentially introduced. The resulting dataset therefore contained periods of time during which the mouse was exposed to a novel juvenile or alone during 4 successive epochs (epochs ‘A’, ‘B’, ‘C’, and ‘D’). In the static context all four epochs occurred in the home cage, whereas in the dynamic context epochs A and B occurred in the home cage, while C and D took place in a distinct environment / context.

**Figure 1:**
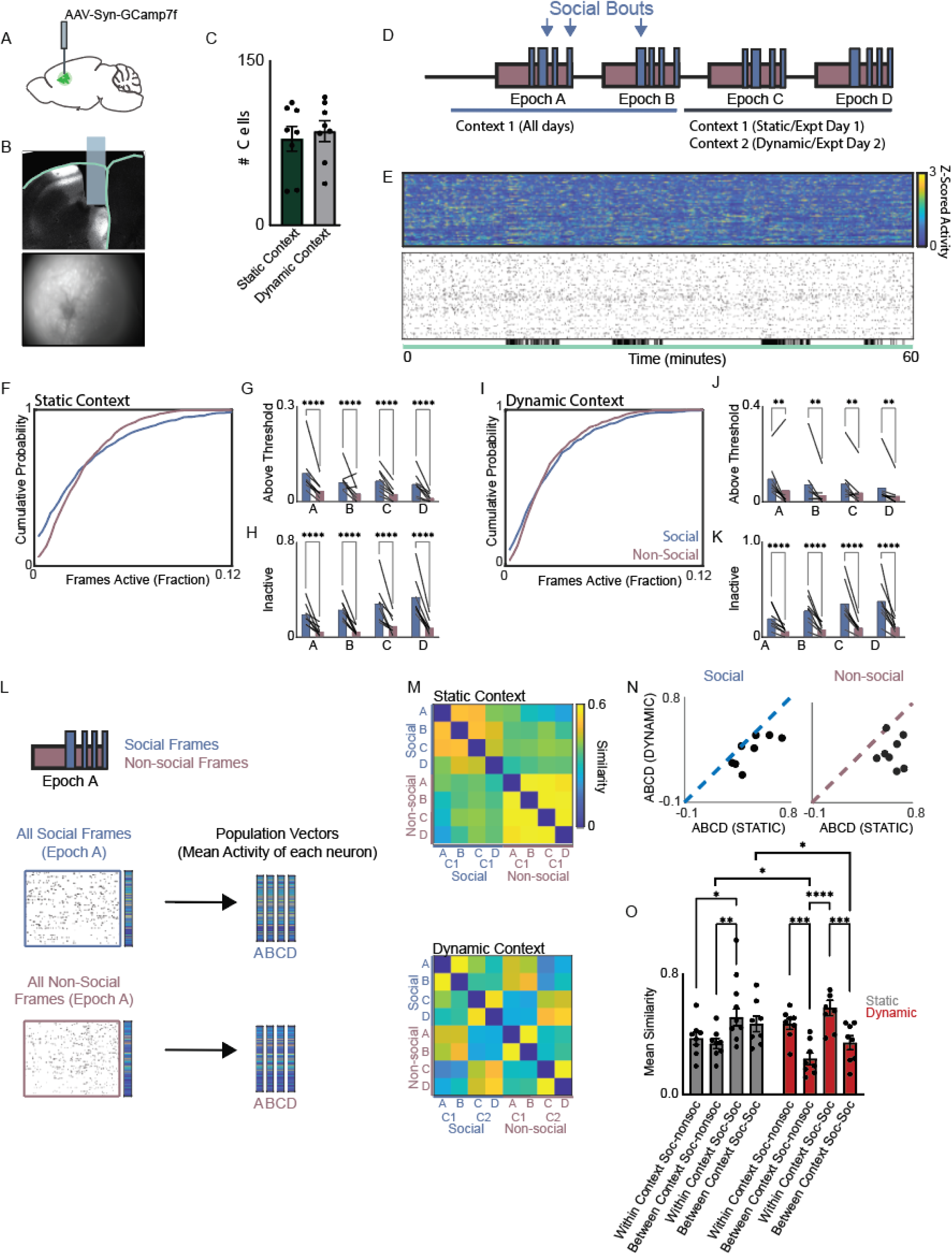
Reorganization of prefrontal network-level activity underlies dynamic contextual experience. **A & B.** AAV8-hSyn-gCamp7f was stereotactically injected into the mPFC of adult mice to visualize neuronal activity via an implanted GRIN lens. B. Coronal section showing GRIN lens placement (top) and an example FOV during behavior. **C.** Bar graph showing the number of cells segmented from Ca^2+^ imaging datasets in each mouse (mean 86 +/- 10 cells for static context vs. 79 +/- 11 cells for dynamic context, n = 8 mice). **D.** Ca^2+^ activity was visualized during serial interactions with 4 mice sequentially introduced to the home cage (static context). On a different day, activity was recorded during two interactions in the home cage followed by two interactions in a second familiar environment (dynamic context). **E.** Example Z-scored Ca^2+^ traces (top) and binary raster of identified events during static context imaging session. Black lines below the raster indicate social interaction bouts. **F.** Cumulative probability function for the proportion of frames that each neuron was active during social bouts (blue) or non-social periods (red) over the entire experiment (left) or during individual epochs (right). Social interaction was associated with an increase in both highly active and inactive neurons in the static context experiment (all epochs social mean 2.5 +/- 0.2% vs. all epochs non-social mean 2.4 +/- 0.3%, p < 0.01, KS Test, n = 688 neurons from 8 mice). **G.** Bar graph showing the proportion of highly active neurons (>7.5% of frames) during social bouts and non- social periods for each epoch in the static context (Epoch A: 9.2 +/- 0.03% in social vs. 3.5 +/- 1.0% in non- social; Epoch B: 6.2 +/- 1.6% in social vs. 2.7 +/- 1.1% in non-social; Epoch C: 6.7 +/- 1.3 in social vs. 2.5 +/- 0.1% in non-social; Epoch D: 5.6 +/- 1.3% in social vs. 1.3 +/- 0.1% in non-social; column factor p < 0.0001, 2- way RM ANOVA with post-hoc Šidák correction, n = 8 mice). **H.** Bar graph showing the proportion of inactive neurons during social interaction and non-social behavior for each epoch in the static context (Epoch A: 19.7 +/- 3.1% social vs. 5.0 +/- 1.5% in non-social; Epoch B: 23.5 +/- 3.1% in social vs. 4.9 +/- 1.3% in non-social; Epoch C: 28.6 +/- 6.1% in social vs. 9.7 +/- 2.8% in non-social; Epoch D: 34.1 +/- 6.1% in social vs. 8.4 +/- 8.4% in non-social; column factor p < 0.0001, 2-way RM ANOVA with post-hoc Šidák correction, n = 8 mice). **I.** Cumulative distributions of activity as in F but for dynamic context (all epochs social mean 2.5 +/- 0.4% vs. non-social mean 2.3 +/- 0.5%; NS by paired t test, p < 0.05, KS test, n = 632 neurons from 8 mice). **J.** Bar graph showing the proportion of highly active neurons (>7.5% of frames) during social interaction and non-social bouts by epoch in the dynamic context (Epoch A: 9.9 +/- 2.7% in social vs. 5.1 +/- 4.3% in non- social; Epoch B: 7.5 +/- 3.8% in social vs. 2.9 +/- 2.0% in non-social; Epoch C: 7.8 +/- 3.3% in social vs. 4.1 +/- 2.4% in non-social; Epoch D: 6.0 +/- 3.0% in social vs. 2.7 +/- 1.7% in non-social; column factor p < 0.0001, 2- way RM ANOVA with post-hoc Šidák correction, n = 8 mice). **K.** Bar graph showing the proportion of inactive neurons during social and non-social bouts by epoch in the dynamic context experiment (Epoch A: 20.1 +/- 4.1% in social vs 6.6 +/- 1.3% in non-social; Epoch B: 28.5 +/- 5.1% in social vs. 8.5 +/- 2.1% in non-social; Epoch C: 35.7 +/- 7.7% in social vs. 10.5 +/- 2.6% in non-social; Epoch D: 38.3 +/- 7.3% in social vs. 11.1 +/- 2.6% in non-social; column factor p < 0.0001, 2-way RM ANOVA with post-hoc Šidák correction, n = 8 mice). **L.** Schematic: we compared network activity during social and non-social bouts within different epochs (A, B, C, or D) by calculating population activity vectors. **M.** Similarity matrices constructed by computing pairwise correlations between population activity vectors generated for social or non-social bouts during each epoch (A-D) within either the static (top) or dynamic context (bottom) experiment. **N.** Scatterplot of the average similarity between population activity vectors associated with social interaction (left) or non-social periods (right) within the static (x-axis) or dynamic (y-axis) context. Each dot is the mean of epochs A vs C, A vs D, B vs C, and B vs D; mean correlations, social: 0.47 +/- 0.05 in static vs 0.34 +/- 0.05 in dynamic, p < 0.01, signed-rank; mean correlations, non-social: 0.60 +/- 0.03 in static vs. 0.31 +/- 0.05 in dynamic, p < 0.05, signed-rank). **O.** Bar graph depecting the mean similarity (correlation) between population activity vectors associated with social interaction or non-social bouts within the same context (i.e., epochs A & B or C & D) or between contexts (i.e., epochs A & C or B & D) in the static (gray bars, left) or dynamic (red bars, right) context experiments (correlation between social interactions, static context: 0.51 +/- 0.06 within-context vs. 0.47 +/- 0.05 between- contexts; dynamic context: 0.58 +/- 0.05 within-context vs. 0.34 +/- 0.05 between contexts; correlation between social and non-social bouts, static context: 0.37 +/- 0.04 within-context vs. 0.34 +/- 0.05 between-contexts; correlation between social and non-social bouts, static context 0.47 +/- 0.03 within-contexts vs. 0.24 +/- 0.04 between-contexts; 2-way ANOVA: p = 0.69 for Experiment Day, p < 0.005 for Comparison Type (social-social and social-nonsocial comparisons within or between context) and p < 0.01 for Experiment Day (static vs. dynamic context) x Comparison Type, posthoc testing performed using Tukey’s multiple comparison test).

We recorded activity from similar numbers of neurons in the static and dynamic context cases (**Figure 1C**; mean 86 +/- 10 cells in the static context, 79 +/- 11 cells in the dynamic context). Individual recordings were segmented using a combined PCA/ICA algorithm (*25*) to identify individual neurons, and events were detected as previously described (*12*) to generate binary rasters (**Figure 1E**) for each mouse corresponding to periods when each neuron was active during the recording.

We next calculated the activity of each neuron during each epoch for periods of social interaction or nonsocial periods, for both the static and dynamic context. ‘Social interaction’ was limited to timepoints when the mouse was actively interacting with the novel juvenile, whereas ‘nonsocial’ included both timepoints when the juvenile was present but the mouse was not engaged in interaction plus the preceding period when the mouse was alone (before the juvenile was introduced). Notably, there was a shift in the shape of the cumulative distribution, such that more cells had either very low or very high activity during social interaction in both the static (**Figure 1F and Supplementary Figure 1**; all social epochs pooled: p < 0.01, K-S Test; individual epochs A-D all p < 0.0001, K-S Test; n = 688 neurons from 8 mice) and dynamic context (**Figure 1I and Supplementary Figure 1**; all social epochs pooled: p < 0.05, K-S test; Epoch A: p < 0.001, K-S test; Epochs B-C, p < 0.0001, K-S test; Epoch D: p < 0.05, K-S test; n = 632 neurons from 8 mice). We obtained similar results when we analyzed data by mouse – again, a significant difference in the number of neurons which were very active (>7.5% of frames) during social interaction (static context: p < 0.0001, 2-way RM ANOVA, n = 8 mice, **Figure 1G**; dynamic context: p < 0.0001, 2-way RM ANOVA, n = 8 mice, **Figure 1J**). There was a similar difference in the number of silent neurons (static context: p < 0.0001, 2-way RM ANOVA, n = 8 mice, **Figure 1H**; dynamic context: p < 0.0001, 2-way RM ANOVA, n = 8 mice; **Figure 1K**).

This shift in the distribution of activity could be driven by distinct sets of neurons in each epoch, or by a single socially-encoding ensemble whose members were consistently modulated by social interaction across epochs. To generate similarity matrices for each experimental day, we first obtained population activity vectors corresponding to the mean activity of each neuron during each epoch of social interaction or nonsocial; then we calculated the similarities, i.e., pairwise correlation coefficients, between these population activity vectors (**Figure 1L-M**). In the static context, population activity vectors for social interaction were very similar to each other, but distinct from those associated with nonsocial periods (**Figure 1M, Top**). By contrast, in the dynamic context, population activity vectors for social interaction and nonsocial activity were similar to each other if they were from the same context, but very different from both types of vectors from the other context (**Figure 1M, Bottom**).

To further quantify these changes in ensemble encoding we compared the similarity between social and nonsocial population activity vectors from epochs A and B with the same type of vectors (social or nonsocial) from epochs C and D. In both the social and nonsocial cases, these similarities were significantly decreased in the dynamic compared to the static context (**Figure 1N**; Social: static context mean correlation coefficient 0.47 +/- 0.05, dynamic context mean correlation coefficient 0.34 +/- 0.05, p < 0.01, signed-rank; Nonsocial: static context mean correlation coefficient 0.60 +/- 0.03, dynamic context mean correlation coefficient 0.31 +/- 0.05, p < 0.05, signed-rank; n = 8 mice). We obtain similar results computing the similarity between population vectors averaged over epochs A and B and those averaged over epochs C and D, as well as when we computed the similarity for vectors from epochs B and C (**Supplementary Figure 2**).

We next quantified the similarity (correlation) between population activity vectors of different types, i.e., social vs. nonsocial (**Figure 1O**). We computed this separately ‘within-context’ (i.e., between epochs A and B and also between C and D) and ‘between-context’ (i.e., between either A or B and either C or D). Note that in the static context condition, the context for epochs A and B vs. C and D will actually be the same. Correspondingly, in the static context condition, the similarity between social and nonsocial activity vectors was not different within vs. between-context (0.37 +/- 0.04 vs. 0.34 +/- 0.05). In the static context condition, the similarity between social activity vectors was also not different within vs. between-context (0.51 +/- 0.06 vs. 0.47 +/- 0.05). However, in the dynamic context condition (in which epochs A and B have a different context than epochs C and D), social and nonsocial patterns of activity were significantly more similar within than between contexts (0.47 +/- 0.03 vs. 0.24 +/- 0.04, p < 0.001). In the dynamic context condition, social activity vectors were also more similar within than between contexts (0.58 +/- 0.05 vs. 0.34 +/- 0.05; p < 0.01; p < 0.01 for Experiment Day x Ensemble Similarity interaction by 2-way ANOVA; posthoc testing performed using Tukey’s multiple comparison test). We obtained similar results (a decrease in the cross-context similarity of social activity vectors) was also still evident when we restricted analysis only to neurons that were active in both contexts (**Supplementary Figure 3**).

Taken together, these data indicate that at the population level, activity patterns observed during social interaction are strongly affected by context. In fact, in the dynamic context condition, social activity vectors were on average more similar to nonsocial activity vectors from the same context (mean correlation of 0.47) than to social activity vectors from the other context (mean correlation of 0.34). This raises questions about whether the encoding of social interactions could be robust to changes in context. To directly examine this, we trained a linear classifier to distinguish social from nonsocial timepoints based on the observed neural activity. Surprisingly, a linear classifier trained on all epochs then tested on held out frames distinguished social vs. nonsocial periods with similar accuracy regardless of the static vs. dynamic context condition (**Figure 2A-B**; mean performance on real data: 68.8 +/- 1.6% in static vs 68.2 +/- 1.7% in dynamic, p = 0.75, signed-rank test; mean performance on shuffled data 56.5 +/- 0.7% in static vs. 55.3 +/- 0.8% in dynamic; n = 8 mice). Note: the classifier performed with similar accuracy when trained and tested within individual epochs for both the static vs. dynamic context condition (**Supplementary Figure 4A-B**).

**Figure 2:**
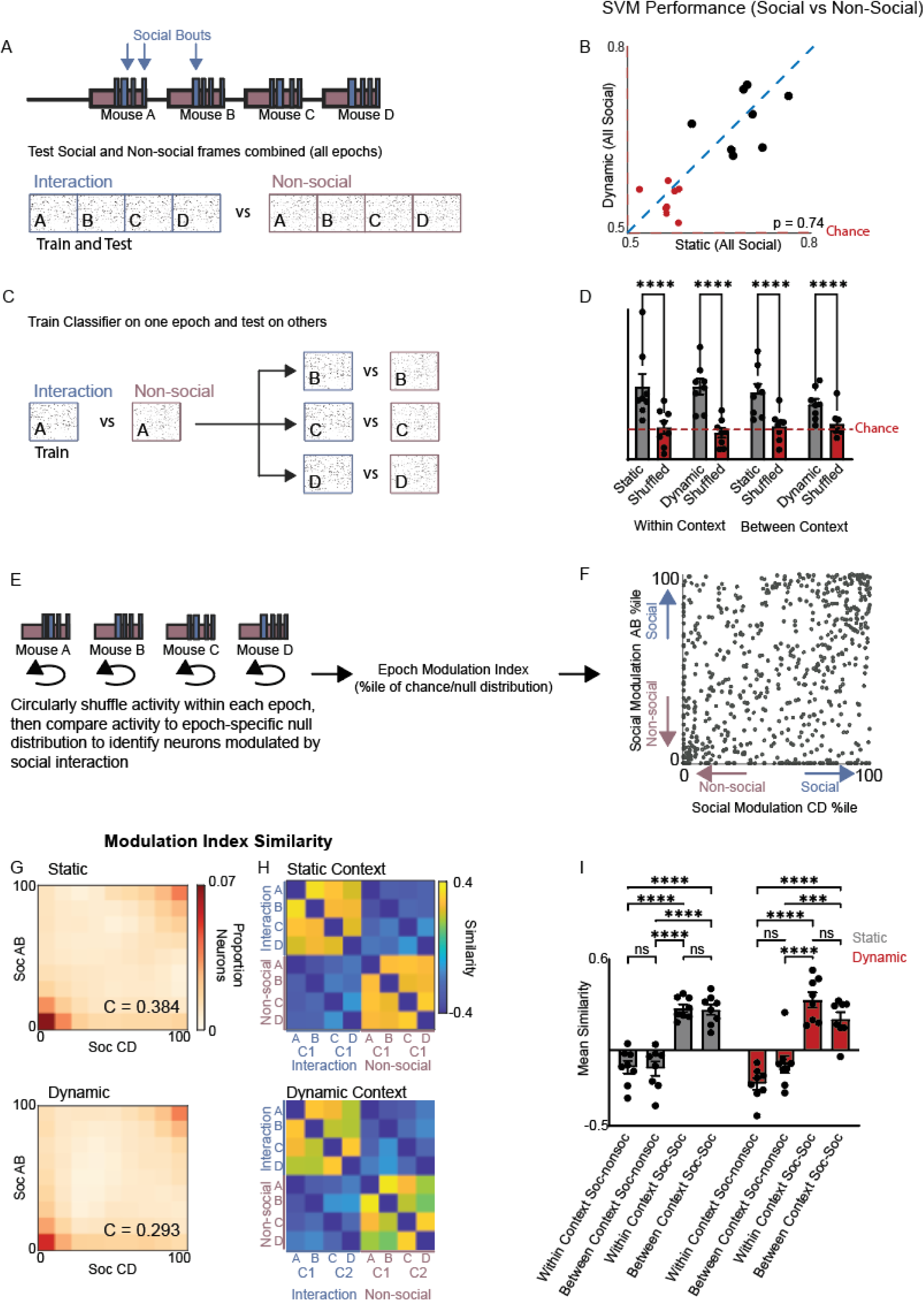
Context-invariant representations of social interactions. **A.** A Support Vector Machine (SVM) classifier was trained using binarized Ca^2+^ activity to discriminate frames corresponding to either social interaction or non-social periods across all epochs. **B.** Performance in static (x-axis) vs. dynamic context (y-axis) (mean % correct: 68.8 +/- 1.6% in static vs. 68.2 +/- 1.7% in dynamic, p = 0.8, signed-rank; shuffled values in red: 56.5 +/- 0.7% for static, 55.3 +/- 0.8% for dynamic, n = 8 mice). **C.** A linear classifier was trained to distinguish frames corresponding to social interactions vs. non-social periods for one epoch (A, B, C, or D), and then tested on the other epochs. **D.** Bar graph of the classifier performance, based on whether the epochs used for training and testing were from the same context (‘within context,’ e.g., A-B or C-D, left) or different contexts (‘between-context,’ e.g., A-C or B-D, right). Classifier performance on real data (gray bars) was compare to the chance level based on randomly shuffled data which maintained the number of active cells in each frame and the overall number of events for each cell (red bars). Classifier performance was similar regardless of experiment day (static vs. dynamic context) or whether within or between-context epoch pairs were used for training and testing (within context: 57.2 +/- 2.1% in static vs. 57.2+/-1.4% in dynamic, between context: 56.2 +/- 1.5% in static vs. 54.3 +/- 0.9% in dynamic; classifier performances on shuffled data: 50.4 +/- 1.1% for static within-context, 49.5 +/- 0.8% for dynamic within-context, 50.6 +/- 0.9% for static between-context and 51.0 +/- 0.8% for dynamic between-context; p < 0.005 for performance on real vs. shuffled data, p = 0.5 for within vs. between context, 2-way RM ANOVA repeating measures by row and column, post-hoc testing using Tukey’s multiple comparisons test). **E.** We calculated the epoch-specific modulation index for each neuron by comparing each its activity during social interactions to a null distribution generated by circularly shuffling its activity across both social and non-social frames within the epoch. **F.** Scatter plot showing the social modulation index for individual neurons calculated across using all social and non-social frames from epochs A&B (y-axis) or Epochs C&D (x-axis). **G.** Heatmaps constructed from 10x10 matrices representing the proportion of neurons that fall into bins corresponding to deciles of social modulation indices. The social modulation index deciles are plotted for epochs A and B on the y-axis and for epochs C and D on the x-axis for the static (top) or dynamic context (bottom) experiments (correlations between social modulation indices in epochs A and B vs. in C and D: static context: 0.38, p < 0.0001; dynamic context: 0.28, p < 0.0001; n = 688 neurons and 632 neurons from 8 mice, respectively). **H.** Similarity matrices constructed by calculating the correlations between vectors of modulation indices for different epochs (A-D) and behaviors (social interaction vs. nonsocial frames) within either the static context (top) or dynamic context (bottom) experiment. **I.** Bar graph summarizing the similarity between vectors of modulation indices for either social interaction (‘Soc)’ or non-social bouts (‘nonsoc’) within the same context (i.e., epochs A & B or C & D) or between contexts (i.e., epochs A & C or D or B & C or D) for either the static (gray bars, left) or dynamic (red bars, right) context experiment (Static context: soc-nonsoc within-context = -0.11 +/- 0.04, soc-nonsoc between context = -0.12 +/- 0.05, within-context soc-soc = 0.28+/-0.03, between-context = 0.26+/-0.03; Dynamic context: soc-nonsoc within-context = -0.22 +/- 0.04, soc-nonsoc between-context = -0.09 +/- 0.06, soc-soc within-context = 0.33+/- 0.05, soc-soc between-context 0.21+/-0.04; 2-way RM ANOVA, p > 0.0001 for comparison (social-social and social-nonsocial comparisons within or between context), p = 0.09 for experiment day (static vs dynamic) p = 0.12 for interaction; posthoc testing performed using Tukey’s multiple comparisons test.

Strikingly, the classifier performed significantly above chance levels even when trained on a single epoch then tested on an epoch from the other context, in both the static and dynamic context conditions (**Figure 2C-D**; mean between-context performance for real data: 56.2 +/- 1.5% in the static condition, 54.3 +/- 0.9% in the dynamic condition; for shuffled data: 50.6 +/- 0.9% in the static condition, 51.0 +/- 0.8% in the dynamic condition; p < 0.0001 for real vs. shuffled in both the static and dynamic conditions by 2-way RM ANOVA; P < 0.005 for main effect of shuffling, p = 0.5 for comparisons across Epoch/experiment day, 2-way RM ANOVA). This indicates that despite the context-dependent reorganization of population-level activity patterns, classifiers, even when trained on a single epoch and single context, learn to identify specific patterns of activity which encode social interaction and are context-invariant.

To better understand the basis for this context-invariant encoding, we sought to determine whether the same neurons consistently encode social interaction across different contexts via positive or negative modulations of their activity. For this, we calculated a ‘social modulation index’ (and ‘nonsocial modulation index’) within each epoch, by circularly shuffling the activity trace of each neuron across all social and nonsocial frames within each epoch to generate a null distribution (**Figure 2E**), then expressing the observed activity level of that neuron as a percentile relative to this null distribution. In this way, neurons that increased or decreased activity during social interaction would have a high (e.g., >90%ile) or low (e.g., <10%ile) social modulation index, respectively. We first calculated a joint social modulation index for epochs A and B (‘AB’), and separately, for epochs C and D (‘CD’) (**Figure 2F**). For both the static and dynamic context conditions, neurons that were strongly positively or negatively modulated by social interaction in AB tended to be similarly modulated in CD, as evidenced by a positive correlation between the vector of social modulation indices in AB and the corresponding vector in CD (**Figure 2G**; static context: AB vs CD correlation coefficient = 0.384, *p* <0.0001; dynamic context: AB vs CD correlation coefficient 0.281, *p* < 0.0001).

To further explore this encoding invariance, we generated similarity matrices by calculating the correlation between vectors of social or nonsocial modulation indices in each epoch (A, B, C, or D) for both the static (**Figure 2H**, top) and dynamic (**Figure 2H**, bottom) context conditions. This showed that social modulation index vectors from each epoch were similar to social modulation index vectors from other epochs, but distinct from nonsocial modulation index vectors (**Figure 2I**). The similarity between social modulation index vectors was significantly greater than between social and nonsocial vectors for both within and between-context comparisons in both the static and dynamic-context conditions.Taken together, these data confirm that despite context-driven changes in population-level activity patterns, the ensemble of neurons whose activity is strongly modulated by social interactions is largely consistent across contexts.

Next, we sought to better understand the relationship between this socially-encoding neuronal ensemble, and context-driven changes in neuronal activity. Specifically, as described earlier, we wondered whether this social ensemble might be non-overlapping and/or orthogonally oriented with respect to representations of contextual information. To test this, we calculated the modulation index vectors based on the degree to which neurons were modulated by context (A-B vs C-D) (**Figure 3A-B**). Specifically, for each neuron, we computed the difference in activity observed during epochs A and B vs. during C and D, and then expressed this as a percentile relative to the distribution of such differences calculated from circularly-shuffled data. We also calculated social (or nonsocial) modulation index vectors for each context (either epochs A and B or C and D) by calculating the average activity of each neuron during social (or nonsocial) frames during epochs A and B (or C and D), and then expressing this as a percentile relative to the distribution observed in circularly-shuffled data (**Figure 3C**). This resulted in a set of modulation vectors for each mouse which represented either the degree to which the activity of each neuron was modulated by context, or the degree to which each neuron’s activity was modulated by social interaction (within a context). Finally, we generated a similarity matrix for these modulation vectors (by computing the correlations between them). Social modulation vectors derived from epochs A and B (‘social AB’) were similar to those derived from epochs C and D (‘social CD’) (**Figure 3D- E**; social AB-social CD correlation: static context 0.37 +/- 0.07 vs. 0.07 +/- 0.03 for shuffled data, p < 0.005; dynamic context 0.27 +/- 0.06 vs. 0.00 +/- 0.06 for shuffled data, p < 0.01). Social AB vectors were also anti- parallel to (i.e., negatively correlated with) nonsocial modulation vectors (‘nonsocial CD’) on both the static and dynamic context days (social AB-nonsocial CD correlation: static context -0.30 +/- 0.09 vs. 0.01 +/- 0.05 for shuffled data, p <0.005; dynamic context -0.20 +/- 0.05 vs. -0.02 +/- 0.04 for shuffled data, p < 0.05). In contrast, on both days the similarity between the social modulation vectors and context modulation vectors was near 0 (indistinguishable from the similarity to shuffled data-derived vectors) indicating that the these representations were near-orthogonal (i.e., the angle between social and context modulation vectors was ∼90 degrees) (social AB-context AB correlation: static context 0.08 +/- 0.04 vs. -0.02 +/- 0.04 for shuffled data, p = 0.50; dynamic context 0.07 +/- 0.04 vs. 0.01 +/- 0.05 for shuffled data, p = 0.83; social AB-context CD correlation: static context -0.07 +/- 0.04 vs. 0.00 +/- 0.06 for shuffled data, p = 0.80; dynamic context -0.06 +/- 0.04 vs. 0.04 +/- 0.03 for shuffled data, p = 0.99; p < 0.0001 for comparison vector type (social vs. nonsocial vs. context), p = 0.56 for day (static vs. dynamic context), p < 0.0001 for interaction, 2-way RM ANOVA).

**Figure 3:**
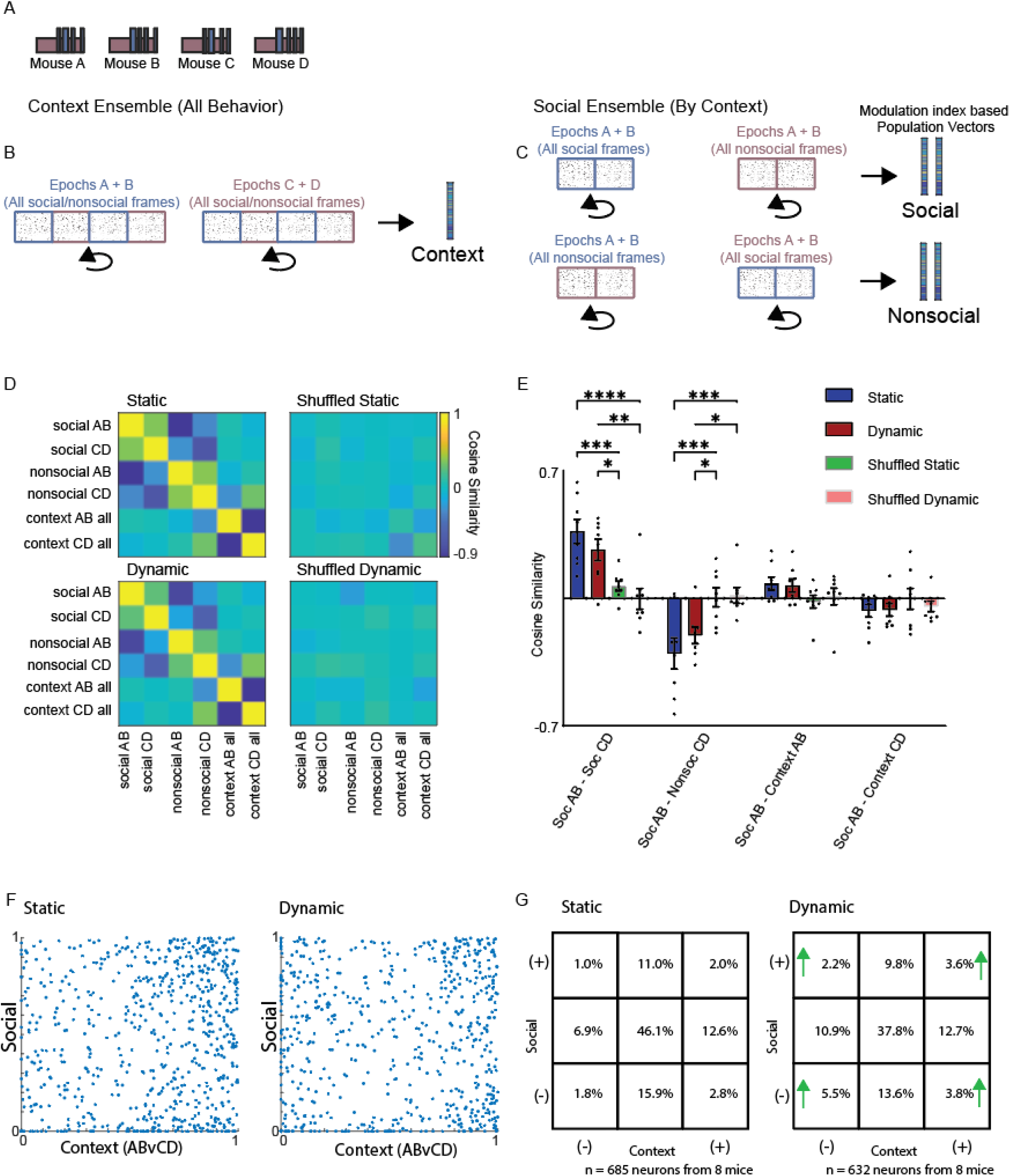
Representations of social information and context are heavily overlapping but orthogonal. **A.** To examine the relationship between ensemble representations of social interaction and context we divided each dataset into sub-rasters corresponding to period of social interaction or non-social behavior (non-social rasters comprised frames that occurred between the social bouts plus the 6000 frames preceding introduction of the novel conspecific). **B.** We generated vectors of modulation indices corresponding to the context-specific modulation of each neuron (i.e., the difference in their activity during epochs A and B vs. C and D, based on both social and non- frames, expressed as a %ile relative to the difference expected by chance based on shuffling each neuron’s activity 10,000 times). **C.** We generated vectors of modulation indices corresponding to social behavior (top) by calculating the difference between each neuron’s activity during social (Soc) vs. nonsocial (Nonsoc) frames within a single context (i.e. either epochs AB or CD). This difference was expressed as a %ile relative to the difference expected by chance based on shuffled data. We generated vectors corresponding to nonsocial modulation indices via an analogous approach (bottom). **D.** We examined the similarity between pairs of social or context-specific ensembles by calculating correlation between vectors of the corresponding modulation indices. This was done for both real (left) and shuffled (right) datasets. **E.** Bar graph of similarities between social, nonsocial, and context-specific modulation index vectors on the static context (blue bars) or dynamic context (red bars) experiment days. Shuffled vectors were generated by randomly re-assigning the neuron associated with each modulation index value. The similarity between context and behavior-specific vectors do not differ from the chance level calculated using shuffled vectors (similarity of Soc AB to Soc CD: 0.37 +/- 0.07 in static context vs. 0.07 +/- 0.03 for shuffled, 0.27 +/- 0.06 in dynamic context vs. 0.00 +/- 0.06 for shuffled; similarity of Soc AB to Nonsoc CD: -0.30 +/- 0.09 in static context vs. 0.01 +/- 0.05 for shuffled, -0.20 +/- 0.05 in dynamic context vs. -0.02 +/- 0.04 for shuffled; Similarity of Soc AB to Context AB: 0.08 +/- 0.04 in static context vs. -0.02 +/- 0.04 for shuffled, 0.07 +/- 0.04 in dynamic context vs. 0.01 +/- 0.05 for shuffled; Similarity of Soc AB to Context CD: -0.07 +/- 0.04 in static context vs. 0.00 +/- 0.06 fo shuffled, -0.06 +/- 0.04 in dynamic context vs. 0.04 +/- 0.03 for shuffled; p = 0.56 for Experiment day (static vs. dynamic context), p < 0.0001 for both comparison (real and shuffled soc-soc, soc-nonsoc, soc-context) and comparison-experiment day interaction, 2-way RM ANOVA (repeating by row and column), posthoc testing performed using Tukey’s multiple comparisons test). **F.** Scatter plots of the social modulation index (calculated in context AB) vs. the context modulation index of each neuron in the static (left) or (dynamic) context experiment. **G.** 9X9 table showing the proportion of neurons which were strongly positively (+) or negatively (-) modulated (i.e., modulation index >90%ile or < 10%ile) by social interaction or context. Green arrows demarcate squares in which greater overlap is observed in the dynamic context compared to static (positive social modulation & positive context modulation p < 0.005, χ^2^ test; positive social & negative context modulation p < 0.0001, χ^2^ test; negative social & positive context modulation p < 0.05, χ^2^ test; negative social & negative context modulation p < 0.0001, χ^2^ test).

Social and context representations could be orthogonal because they are composed of non-overlapping neuronal ensembles. Alternatively, neurons which are strongly positively modulated by social interaction could be evenly divided between positive and negative context-modulation (and the same for neurons which are strongly negatively modulated by social interaction). Because our experiment included days with both static and dynamic contexts, we could compare overlap between social and context representations (the amount of overlap observed on the dynamic context day) to the level expected by chance / the passage of time (the amount of overlap on the static context day). Specifically, we could organize cells which were strongly positively (>90%ile) modulated, non-modulated (10-90%ile) or negatively modulated (<10%ile) by social interaction or context into a 3x3 table (**Figure 3F-G),** construct a different table for the static vs. dynamic context day, and determine whether any of the strongly modulated groups (positive social-positive context; positive social-negative context; negative social-positive context; negative social-negative context) overlap more on the dynamic than static day. Surprisingly, we observed significantly more overlap on the dynamic day compared to the static day in every case (positive social-positive context: 3.6% in dynamic vs. 2.0% in static, p < 0.01; positive social-negative context: 2.2% in dynamic vs. 1.0% in static, p < 0.0001; negative social-positive context: 3.8% in dynamic vs. 2.8% in static, p < 0.05; negative social-negative context: 5.5% in dynamic vs. 1.8% in static, p < 0.0001; χ^2^ test; n = 632 cells in dynamic and 685 cells in static). Thus, there is actually greater-than-expected overlap between social and context ensembles, however this overlap includes neurons modulated in opposing directions, maintaining an approximately orthogonal relationship between these representations.

Finally, we tested whether these general principles (orthogonality of representations enabling generalization across contexts) extend to the role of the prefrontal cortex in a distinct behavior, namely anxiety-related avoidance. For this, we measured prefrontal neuronal activity in mice over multiple epochs in which they explored an elevated zero maze (EZM), elevated plus maze (EPM), and a T-shaped maze (T-Maze) (**Figure 4A**). The EZM and EPM both utilize open and closed arms positioned atop an elevated platform and are commonly used to assay anxiety-related avoidance in rodents (rodents tend to avoid the exposed open arms and prefer the safe closed arms). By contrast, the T-Maze consists of three closed arms (two long arms + one short arm). Each epoch was preceded and followed by a period of recording when the mouse was alone in its home cage to permit the identification of activity patterns specifically associated with each maze. Epochs were also interleaved with two 20-minute rest periods during which the camera was turned off.

**Figure 4.**
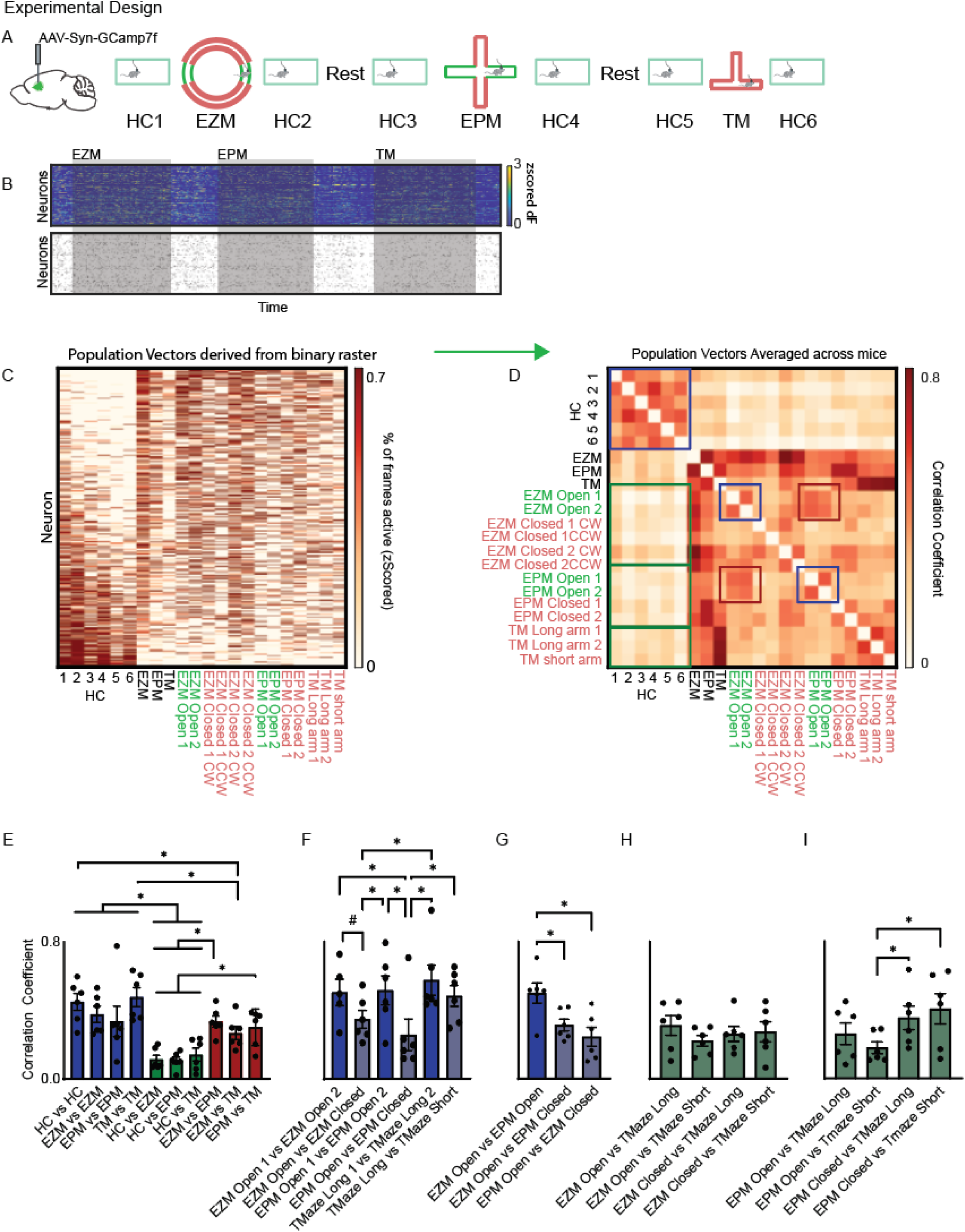
Context-invariant patterns of activity for anxiety-related information. **A.** Experimental design: Ca^2+^ activity was recorded throughout a session in which mice explored the elevated zero maze (EZM), elevated plus maze (EPM), and a T-shaped mazeI). Mice were recorded within their home cage before and after exploring each of these novel mazes, and recording blocks were separated by 30 minutes of rest. **B.** Example Z-scored Ca^2+^ activity (top) and binary activity raster (bottom). We recorded activity from a total of 633 neurons in 6 mice. **C.** Behavior was subdivided into each home cage epoch and the time mice spent within each arm of the EZM, EPM, and T-Maze. EZM closed arms were further divided by the direction of travel (clockwise, CW; counter- clockwise, CCW). We generated population activity vectors for each epoch and arm by calculating the mean activity of each neuron within each maze and each subregion (arm). Neurons were concatenated across all mice, then sorted based on average activity during home cage epochs, revealing different sets of neurons active in the home cage vs. during exploration of other environments. **D.** We generated a similarity matrix by computing correlations between population activity vectors corresponding to different epochs / specific arms for each mouse, then averaging the resulting 6 similarity matrices. **E.** Bar graph of average similarities (correlations) between population activity vectors from various mazes. HC vs. HC = average similarity of population activity vectors from adjacent home cage epochs, e.g., HC1 vs. HC2. EZM vs. EZM = average similarity of all pairs of different population activity vectors from the EZM, e.g., EZM closed arm 1 CW vs. EZM open arm 2, etc. EPM vs. EPM and TM vs. TM were computed similarly. Data in E-I are subsets of mixed-effects analysis with matching across rows and between column p-value < 0.0001, post- hoc control of false discovery rate performed using two-stage method of Benjamini, Krieger, and Yekutieli, n = 6 mice except for EPM Open vs EPM Open comparison as one mouse did not explore one of the arms. **F.** Bar graph of the averaged similarities (correlations) between population activity vectors associated with specific arms within the same maze. #: q value = 0.059. **G-I.** Bar graphs of the averaged similarities (correlations) between population activity vectors associated with specific arms in different mazes.

We recorded 633 neurons from 6 mice. We segmented individual neurons and converted the activity of each neuron to a binary raster corresponding to frames in which each neuron was active or inactive (**Figure 4B**). We next examined population activity vectors pooled across mice corresponding to the mean activity of each neuron either during each home cage epoch, within each maze, or within specific arms of a maze (**Figure 4C + Supplemental Figure 5**). Visual examination of pooled population activity vectors showing activity across all mice during specific epochs in the home cage or each maze illustrated gross differences in network activity related to changes in context. We then generated population activity vectors for specific types of exploration (home cage, EZM / EPM / TMaze, EZM / EPM open / closed arms, etc.) and each mouse, calculated correlations between these vectors to generate similarity matrices (for each mouse), then averaged across mice to obtain a mean similarity matrix (**Figure 4D**). This similarity matrix shows that distinct patterns of activity are associated with home cage exploration vs. the different mazes (**green boxes in Figure 4D and Supplementary Figure 5**). Specifically, the similarity between activity vectors associated with different arms of the same maze was significantly higher than the similarity between activity vectors for each maze and those associated with home cage exploration (**blue vs. green bars in Figure 4E**; EZM vs EZM mean correlation coefficient 0.38+/-0.05, EPM vs EPM mean correlation coefficient 0.34+/-0.09, TMaze vs Tmaze mean correlation coefficient 0.48+/-0.05, HC vs EZM mean correlation coefficient 0.12+/-0.02, HC vs EPM mean correlation coefficient 0.11+/-0.02, HC vs Tmaze mean correlation coefficient 0.14+/-0.02; mixed-effects analysis, p < 0.0001). Furthermore, the activity vectors for the different mazes were significantly more similar to each other than to home cage activity vectors (**red vs. green bars in Figure 4E**).

As expected, there is high similarity between population activity vectors associated with each of the two open arms from the same maze (EZM or EPM) (**blue boxes in Figure 4D and Supplementary Figure 5**). Specifically, activity vectors for each open arm were significantly more similar to those for the other open arm in the same maze than to activity vectors for the closed arms in the same maze (**Figure 4F**; EZM Open 1 vs EZM Open 2 mean correlation coefficient 0.51+/-0.08, EZM Open vs EPM Closed mean correlation coefficient 0.35+/-0.05, EPM Open 1 vs EPM Open 2 mean correlation coefficient 0.53+/-0.09, EPM Open vs EPM Closed mean correlation coefficient 0.26+/-0.09). However, we also observed very high similarity between population activity vectors associated with the EZM open arms and those associated with the EPM open arms (**red boxes in Figure 4D and Supplementary Figure 5**). Indeed, activity vectors for the EZM open arms were significantly more similar to those for the EPM open arms than those for the EPM closed arms (and vice-versa) (**Figure 4G**; EZM Open vs EPM Open mean correlation coefficient 0.50+/-0.06, EZM Open vs EPM Closed mean correlation coefficient 0.31+/-0.04, EPM Open vs EZM Closed mean correlation coefficient 0.25+/-0.06). Otherwise, activity vectors for different maze subregions were relatively dissimilar, except for the EPM closed arms and the TM long arms, perhaps reflecting geometric similarity (**Figure 4H-I**). We observed similar results using population activity vectors based on Z-scored calcium traces instead of binary activity rasters (**Supplementary Figure 6**).

These findings suggest that prefrontal representations of anxiety-related information, like social information, persist across different contexts. We therefore sought to understand the degree to which anxiety-related representations overlap with representations of context. To do this we again generated vectors of modulation indices, corresponding to the degree to which each neuron’s activity differed between contexts or sub-regions of a maze (**Figure 5A-D**). Specifically, we calculated the difference between each neuron’s mean activity in two conditions (e.g., home cage vs. EZM, EZM open vs. closed arms, etc.), then expressed these differences as percentiles relative to distributions based on shuffled data. We did this to identify ensembles of neurons which were differentially active between different home cage epochs (**Figure 5B**), ensembles which distinguished each maze from the adjacent home cage epochs (**Figure 5C**), and ensembles which discriminated different compartments within a maze, e.g., the open and closed arms (**Figure 5D**). We then computed the similarity between different vectors of modulation indices to quantify the similarity between neuronal ensembles encoding specific contexts and/or anxiety-related information (**Figure 5E**). The environment was obviously identical across home cage epochs; thus the similarity between modulation index vectors associated with different pairs of home cage epochs represents a benchmark for evaluating differences in neuronal ensembles expected simply due to the passage of time and accumulation of behavioral experiences over the course of the recording.

**Figure 5.**
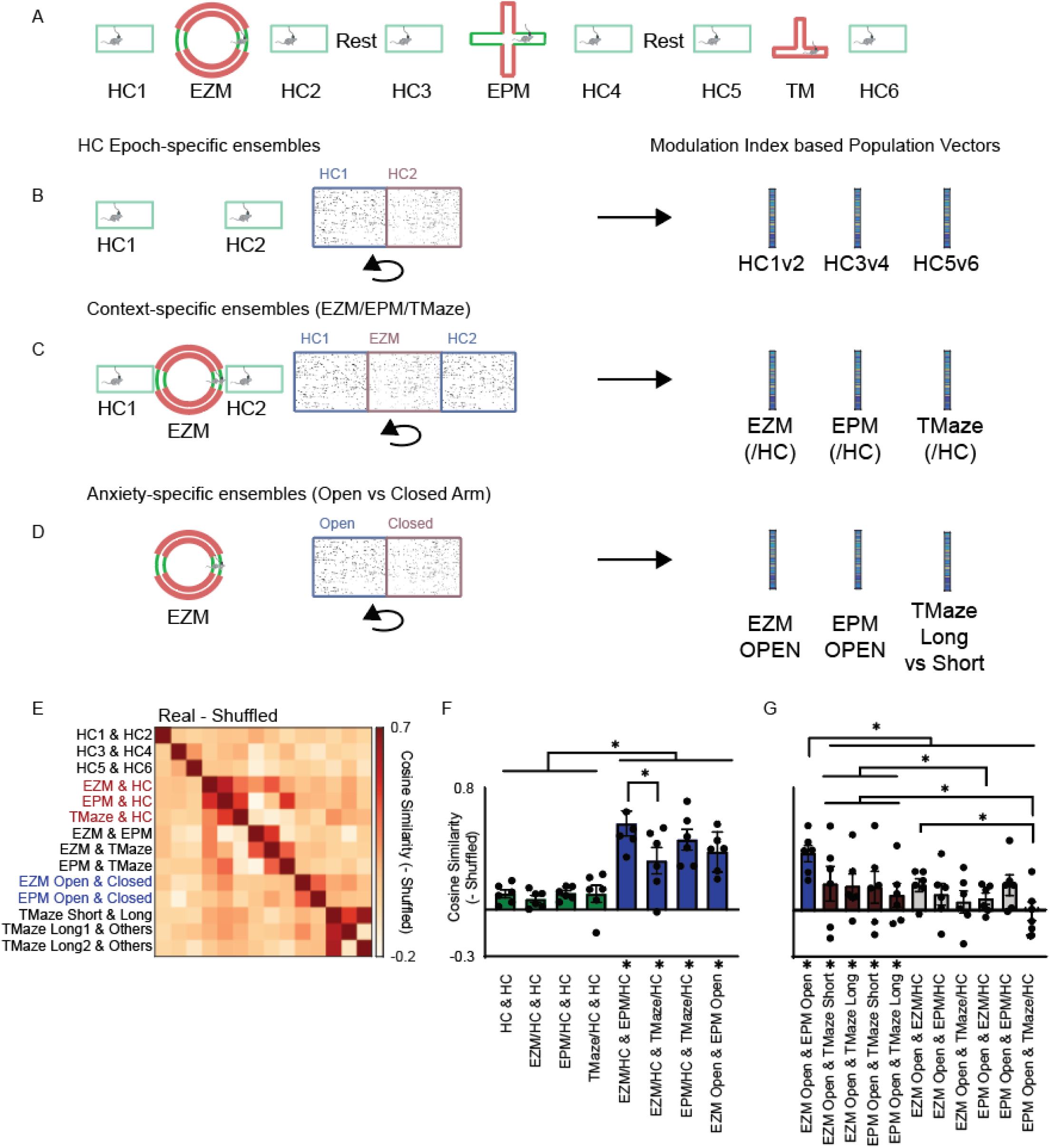
Anxiety-related information and context are encoded by orthogonal ensembles. **A-D.** Modulation index vectors were generated based on activity differences for each neuron between different home cage (HC) epochs (B), between each maze (EZM, EPM, or T-maze) and the adjacent home cage epochs (C), or between different regions within a maze (the open vs closed arms of the EZM or EPM (D), to enable comparisons of context-specific and anxiety-specific ensembles. To calculate the home cage modulation index for a neuron, we expressed its mean activity during one home cage epoch as a %ile relative to a distribution of mean activity values generated by shuffling its activity across both home cage epochs. Context-specific modulation indices for the EZM, EPM, or T-maze were each generated by expressing the mean activity of each neuron within that maze as a %ile relative to a distribution of mean values expected based on shuffling across all frames from that maze and the adjacent HC epochs. Anxiety-specific ensembles were generated by expressing the mean activity of each neuron during open arm exploration as a %ile relative to a distribution of mean values derived after shuffling across all frames from that maze (EZM or EPM). We calculated modulation index vectors for the long or short arms of the T-maze analogously, by comparing mean activity in that arm to a distribution expected by chance derived by shuffling across all T-maze frames. **E.** 14x14 similarity matrices showing the similarity (correlation) between each pair of modulation index vectors (associated with context, anxiety-specific encoding, or specific arms of the T-maze) for real (top) or shuffled (bottom) data. **F.** Bar graph depicting the average similarity (correlation) between pairs of modulation index vectors corresponding to differences between HC epochs (‘HC & HC’), differences between specific mazes and the adjacent HC epochs (EZM/HC, EPM/HC, or TMaze/HC), or anxiety-related information in the EPM vs. EZM (EZM open & EPM open). The similarity between pairs of shuffled vectors (in which modulation index values were randomly re-assigned to different neurons) was near zero. Modulation index vectors for different mazes corresponding to the modulation of neurons within specific contexts were significantly more similar than chance (notably this was not the case for pairs of modulation index vectors corresponding to differences between HC epochs) (P < 0.005 for real vs. shuffled, p < 0.0001 for comparison (HC & HC, EZM/HC & HC, EZM/HC &EPM/HC, etc) and interaction, 2 Way RM ANOVA (repeating by row and column). Panels F and G show subsets of the data from a single statistical analysis, and have been separated into two panels for clarity. Post-hoc control of false discovery rate performed using two-stage method of Benjamini, Krieger, and Yekutiel. n = 6 mice. Shuffled data not shown but * below x-axis indicate similarities that are significantly different (p < 0.05) from those calculated from shuffled values. **G.** Bar graph depicting the average similarity (correlation) between vectors of modulation indices associated with specific arms within a maze, i.e., the open arms of the EZM or EPM, or the long or short arms of the T-maze. n = 6 mice. Shuffled data not shown but * below x-axis indicate comparisons that differ significantly from shuffled data.

The modulation Index vectors distinguishing each maze from adjacent home cage epochs were significantly more similar to each other than were vectors distinguishing different home cage epochs or shuffled data (**green vs. blue bars in Figure 5F**; P < 0.005 for surrogate vs real data, p < 0.0001 for both Ensemble comparison and interaction, 2 Way RM ANOVA). This indicates that largely overlapping ensembles encode the contexts associated with the EZM, EPM, and TMaze. The similarity between modulation index vectors for the EZM open vs. closed arms and vectors for the EPM open vs. closed arms was also significantly higher than the similarity between modulation index vectors for different home cage epochs (**Figure 5F**). This confirms that neuronal ensembles encoding anxiety-related information persist across different contexts. Furthermore, modulation index vectors for the EZM open vs. closed arms and those for the EPM open vs. closed arms were significantly more similar to each other than either was to modulation index vectors distinguishing each maze from home cage epochs (mean similarity between EZM/EPM/TMaze vs. home cage (‘context vectors’) and EPM/EZM open vs. closed (‘anxiety vectors’): 0.12+/-0.03, mean similarity between different anxiety vectors: 0.42 +/- 0.08; mean similarity between different context vectors: 0.44 +/- 0.06, RM ANOVA P < 0.0001, posthoc Tukey’s multiple comparisons test). We observed also similar results utilizing vectors generated from Z-scored calcium traces instead of binary activity rasters (**Supplementary Figure 7**).

Finally, to further assess the context-invariance of anxiety-related representations, and elucidate relationships between context and anxiety-related encoding, we trained support vector machines (SVMs) to distinguish frames corresponding to different contexts (e.g., EZM vs. adjacent home cage epochs), home cage epochs (e.g., home cage epoch 1 vs. 2), or subregions within the same maze (e.g., the open vs closed arms of the EZM). We tested each of these SVMs on each of the 14 pairs of mazes (e.g., EZM vs. HC, EZM vs. EPM, etc.), epochs (HC1 vs. HC2, etc.), and subregions (e.g., EZM/EPM open vs. closed, TM short vs. long) (**Supplementary Figure 8A**). If similar neuronal ensembles encode contexts corresponding to each maze, or anxiety-related information across two contexts (EZM and EPM), then the classifier should perform above chance in one context after having been trained on neural activity from the other. Indeed, we observed that classifier performance largely mirrored the similarities we previously observed for population activity vectors (Figure 5) and modulation index vectors (Figure 6). Specifically, the classifier performed well above chance when trained to distinguish one maze from adjacent home cage frames and then tested on another maze (**Supplementary Figure 8B-C**). The classifier also performed equally well at distinguishing open arm frames from closed arm frames, when trained on one maze (EZM or EPM) and then tested on the other one (**Supplementary Figure 8D**; EZM Open vs. EPM Open mean performance 0.66+/-0.05 (shuffled 0.51+/-0.01), EZM Open vs. Tmaze Short mean performance 0.53+/-0.03 (shuffled 0.51+/-0.01), EZM Open v Tmaze Long mean performance 0.54+/-0.01 (shuffled 0.50+/-0.01), EPM Open v Tmaze Short mean performance 0.52+/-0.02 (shuffled 0.49+/-0.01),. In contrast, SVMs performed at chance levels when they were trained to distinguish context (e.g., EZM vs. HC), and then tested on discrimination of open arms vs. closed arms of the EZM or EPM, again supporting the finding that ensemble representations for context are orthogonal to those for anxiety-related information (**Supplementary Figure 8D;** EZM Open vs. EZM/HC 0.52+/-0.01 (shuffled 0.51+/-0.01), EZM Open vs. EPM/HC mean performance 0.51+/-0.02 (shuffled 0.50+/-0.01), EZM open vs. Tmaze mean performance 0.48+/-0.01 (shuffled 0.51+/-0.01), EPM Open vs. EZM/HC mean performance 0.52+/-0.02 (shuffled 0.50+/-0.01), EPM Open vs. EPM/HC mean performance 0.51+/-0.01 (shuffled 0.50+/- 0.01), EPM Open vs. Tmaze/HC mean performance 0.48+/-0.01 (shuffled 0.49+/-0.01); p < 0.0001 for classifier type, p < 0.01 for real vs. shuffled data, and p < 0.0001 for interaction; 2-way RM ANOVA).

## Discussion

Successful behavior requires reliable representations of pertinent information across varying contexts and conditions. While ensemble representations and other forms of multineuron encoding may increase reliability (*12, 26–29*), information pertaining to multiple behavioral variables will be represented via overlapping ensembles, potentially changing and/or confounding the meaning of changes in the activity of individual neurons. Neuronal activity can also change with the passage of time and accumulation of behavioral experiences (*30, 31*), potentially further interfering with the stability of representations for particular types of information. Numerous studies have quantified the ability of single neurons and multineuron ensembles to encode social information (*10–12, 32–34*). Previous studies have found that social ensembles are orthogonal to those associated with exploratory behavior in the amygdala and distinct from those associated with feeding behavior in orbitofrontal cortex (*11, 32*). Still, it remains unclear how ensembles encoding social information interact with the contemporaneous encoding of other behavioral variables, particularly context, in prefrontal cortex. This is a critical question as contextual features of an environment are known to alter neuronal activity underlying learned behaviors (*23, 35*), and contextual representations within the mPFC play a key role in episodic memory formation and spatial processing (*36*).

Here, we show that neuronal ensembles modulated by social behavior remain largely invariant during changes in the environment, allowing social encoding to generalize across contexts. Notably, we still found extensive context-dependent reorganization of population-level activity *during* social interaction (when we examined population activity vectors), and this context-invariance was only evident when we specifically examined the encoding of specific behavioral variables by individual neurons (using modulation indices) or the entire network (using SVM classifiers). Surprisingly, this context-invariance occurs despite greater than expected overlap between context and social encoding neurons, because these two representations are largely orthogonal. Social ensembles were also orthogonal to ensembles associated with the passage of time in the static context experiment (i.e., ‘drift’) (*31, 37, 38*). Similarly, anxiety-related ensembles, recruited during exploration of maze arms typically viewed as anxiogenic, persist across different maze geometries, enabling context-invariant encoding of anxiety-related information. This type of generalization fits with traditional views of the prefrontal cortex as an ‘executive controller’ that represents abstract features of the environment and behavioral states in order to facilitate decision-making (*39*). Of course, it is unclear whether the context-invariance of these representations arises within the prefrontal cortex or upstream regions, or via interactions between them.

The ability of the prefrontal cortex to multiplex socioemotional and contextual representations using overlapping neurons also aligns with the view of prefrontal neurons as possessing mixed selectivity. Such mixed selectivity could facilitate the formation of multimodal representations that store flexible context-specific associations and/or guide context-specific behaviors. It is unknown whether specific mechanisms or circuit parameters are necessary to achieve this multiplexing and maintain discrete representations in orthogonal configuration. Furthermore, abnormalities in social and anxiety-related behaviors are common manifestations of many neurodevelopmental (*1, 9, 12, 16, 19, 40, 41*) and neurodegenerative disorders (*42–45*). As many studies have begun studying representations of social and anxiety-related information in mouse models of disease, it will be important to explore whether the context-dependence of these representations might be disturbed, leading, for example, to abnormal instability of social encoding or the inappropriate generalization of anxiety-related representations.

## Methods

All experiments were conducted according to the National Institutes of Health (NIH) guidelines for animal research, and protocols were reviewed and approved by the Institutional Animal Care and Use Committee (IACUC) at the University of California, San Francisco (UCSF; protocol number AN185374).

### Stereotactic Injection and Lens Implantation

C57/B6J mice were obtained from Jackson Laboratories (Bar Harbor, ME). We utilized adult mice of either sex housed and bred in the UCSF animal facility. To image prefrontal ensemble activity during freely moving behavior, mice were injected with AAV8.Syn.gCaMP7f.WPRE.SV40 (Penn Viral Core). Coordinates for injection into mPFC were (in mm relative to Bregma) [+1.7 (AP), –0.3 (ML), and –2.3 (DV)], and 1 mm GRIN lenses with an integrated baseplate (Inscopix, Palo Alto) were implanted at the same AP and ML coordinates to a depth of 2.1 mm. Imaging experiments were performed by mounting a microendoscope (Inscopix nVoke) at least 4 weeks after lens implantation to permit recovery.

### Social Behavior

Adult mice were accustomed to the room and handling by observer for at least 3 days prior to experiments, with microendoscope attached. In addition, to mitigate handling or context-induced anxiety, each mouse was habituated to the second context which contained clean bedding for 1 hour daily on 5 consecutive days prior to imaging experiments. On test day, mice were habituated with the scope turned on for 5 minutes prior to starting recordings/acquiring images. During social epochs, novel sex-matched juvenile mice (<6 weeks) were sequentially introduced to the cage containing the test mouse (either the test mouse’s home cage or the second context). Dynamic and static context recordings were obtained on different days and consisted of 10 minute baseline recordings preceding the first and third epoch followed by sequential epochs consisting of 5 minutes of social interaction and interleaved periods in which the mouse was alone. Social behavior was scored manually.

### Anxiety-Related Behavior

Adult mice were habituated to the room and observer for at least 3 days prior to recording. Imaging experiments were performed over a single day with the microendoscope attached throughout the experiment. Imaging experiments consisted of three primary epochs defined by a primary maze which (EZM, EPM, and tMaze) separated by a rest period of 20 minutes during which the microendoscope was turned off. Each epoch consisted of 10-minute recordings in the home cage before and after 30 minutes in which the mouse explored the respective maze.

### Image Acquisition and Segmentation

Images were acquired using an nVoke microendoscope (Inscopix, Palo Alto) attached to a laptop computer and synced to a separate video acquisition computer running Anymaze. Frame rate was 20 Hz, and the light power was 0.2 mW. Acquisition was performed using 4 × 4 pixel binning. We segmented images as previously described using a modified PCA/ICA approach(*12, 25*). Specifically, we used the output from the PCA/ICA to identify a set of contiguous pixels representing a single neuron, then averaging the signal within those pixels. We then subtract the signal within the neuron from the average signal in the surrounding pixels to deconvolve the activity of each neuron from the surrounding neuropil, then low pass filtered the resulting traces to remove high-frequency noise using the MATLAB command: designfilt (“lowpassfir,” “PassbandFrequency,” 0.5, “StopbandFrequency,” .65, “PassbandRipple,” 1, “StopbandAttenuation,” 25). We identified calcium transients based on the amplitude and first derivative of deflections from baseline as previously described(*12*), adjusting parameters corresponding to the first derivative of the dF/F_0_, the amplitude, and the integrated area under the curve of identified events to achieve >95% sensitivity and specificity for each dataset.

### Quantifying ensemble activity

We then converted datasets to binary rasters denoting periods in which each neuron was active over the course of the experiment. We generated population vectors corresponding to the fraction of frames each neuron was active during the specified behavior/epoch. We performed this analysis first by pooling all neurons from each mouse to generate a single population vector for each epoch/behavior. Similarity matrices were generated by calculating the pairwise correlation coefficient between each set of population vectors. Analysis of individual mice was performed by generating population vectors corresponding to the fraction of frames each neuron was active within each behavioral epoch and generating a similarity matrix corresponding to the pairwise correlations between each behavior-specific population vector for each mouse. Population vectors based on dF/F_0_ were generated by calculating the mean Z-scored calcium activity for each neuron within each behavior/epoch from which corresponding similarity matrices were generated as above. We achieved similar results using this approach (**Supplementary Figures**).

### Quantifying behaviorally modulated neurons

The activity of each neuron was quantified as the fraction of frames in which it was active during a particular behavioral epoch. To disambiguate changes in activity driven by social interaction from context-driven changes we calculated the mean activity for each neuron over all frames corresponding to a given behavioral variable in real or shuffled data within a given epoch. We generated a null distribution for each neuron within each epoch by circularly shuffling the data 10,000 times to calculate the activity that would be expected by chance during a given epoch-specific behavior. This permitted us to generate an epoch-specific modulation index for each neuron corresponding to its mean activity during a specified behavior as a percentile relative to the null distribution, with high percentiles (>90^th^ percentile) denoting that the real activity of the neuron during a behavior was greater than expected by chance, and a low percentile (<10 percentile) denoting neurons which were less active during a specified behavior than would be expected by chance (compared to all activity during that epoch). Modulation index vectors contained the modulation index of each neuron within a given behavior/epoch and were used to generate similarity matrices. For the social/context experiment, an epoch was defined as beginning 5 minutes preceding the onset of social interaction with each mouse and ending when a given conspecific was removed from the home cage. For the anxiety/context experiment, an epoch was defined as the entire time that a mouse spent within a given maze.

### Linear Classifier

We trained a Support Vector Machine to distinguish frames corresponding to social interaction from frames corresponding to non-social behavior. Models were trained using the command “fitclinear” in MATLAB. The classifier was first trained and tested on all social and non-social frames from the four epochs and performance was averaged over 500 iterations, leaving 25% hold out. In a separate analysis we trained and tested the classifier on social and non-social frames within each epoch independently, in this case leaving 20% hold out. As the number of frames in which mice engaged in social interaction varied and were not equal to the number of frames in which mice engaged in non-social behavior, we subdivided the data by behavior and randomly selected an equivalent number of frames corresponding to social interaction or non-social behavioron each iteration. Finally, to understand if signatures of neural activity identified in one epoch could inform the classifier across different epochs, we trained the classifier on 100% of frames from each epoch (prior to randomly downsampling on each iteration to balance the data), and then tested the classifier on data from the remaining three epochs. This was performed over 500 iterations. Shuffled data was generated by randomly swapping the identity of the neuron assigned to individual calcium events so that both the number of events in each frame and the number of events assigned to a given neuron was unchanged in the resulting surrogate dataset. We utilized an analogous approach to distinguish frames underlying different mazes, or locations within anxiety-provoking environments. Here we again trained the classifier to distinguish between a pair of contexts or locations based on calcium data, then utilized the trained model to distinguish between a different pair of contexts over 200 iterations. As with social data we utilized 100% of frames within each epoch and utilized balanced training data.

### Orthogonal Ensembles

To compare social ensembles to context specific ensembles we first generated population vectors corresponding to the activity of each neuron during all of the frames in which mice were engaged in social interaction, and all of the frames in which mice were not engaged in social interaction (non-social frames). We then determined a modulation index for each neuron by comparing its activity during social frames to non-social frames. We removed neurons which were not active during either set of frames. We next generated population vectors corresponding to context by calculating the mean activity of each neuron in epochs A&B, and for epochs C&D. We next determined the degree to which each neuron was modulated in epochs A&B compared to C&D using all frames (social + non-social). We then zero-centered the modulation index of each neuron and compared the cosine similarity between resulting social- and context-specific vectors, where:

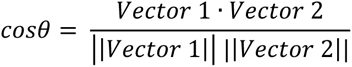

Shuffled comparisons were generated by shuffling cell labels on modulation vectors. We used an analogous approach to compare context-specific and anxiety-specific population vectors as detailed in figure 5.

### Statistical Analysis

Neurons were pooled into population vectors corresponding to entire datasets and activity of these pooled vectors was compared using KS-test and *t*-test as denoted in the manuscript. We then constructed pseudo-ensembles for each behavior corresponding to the activity (or modulation index) of all of the neurons during a specified behavioral epoch. We compared these population vectors by calculating the Pearson correlation coefficient between each pair of vectors. We performed analogous comparisons on population activity and vectors generated for each mouse during each behavior. The activity of population vectors generated for each mouse was compared using repeated measures ANOVA for epoch and behavior (social vs non-social behavior). All ANOVAs were performed using Graphpad Prism. All other statistical comparisons were performed using MATLAB. To compare the degree to which population vectors changed across contexts, we calculated the Pearson correlation coefficient between specified pairs of vectors and compared the result using Wilcoxon signed-rank test. Classifier performance is defined as the mean fraction of frames classified correctly over all iterations in which the classifier was trained and tested. Post-hoc comparisons of population vector correlations, cosine similarity, and classifier performance utilized the two-stage method of Benjamini, Krieger, and Yekutiel to control the rate of false discovery.

## Acknowledgements

This study was funded by the National Institute of Neurological Disorders and Stroke (K08 NS105938-01A1 to NAF) and the National Institute of Mental Health (R01MH117961 to VS), as well as the UCSF Dolby Family Center for Mood Disorders. We thank Kathleen Cho and Damhyeon Kwak for thoughtful comments on the manuscript.

## Supplementary Information

**Supplementary Figure 1:**
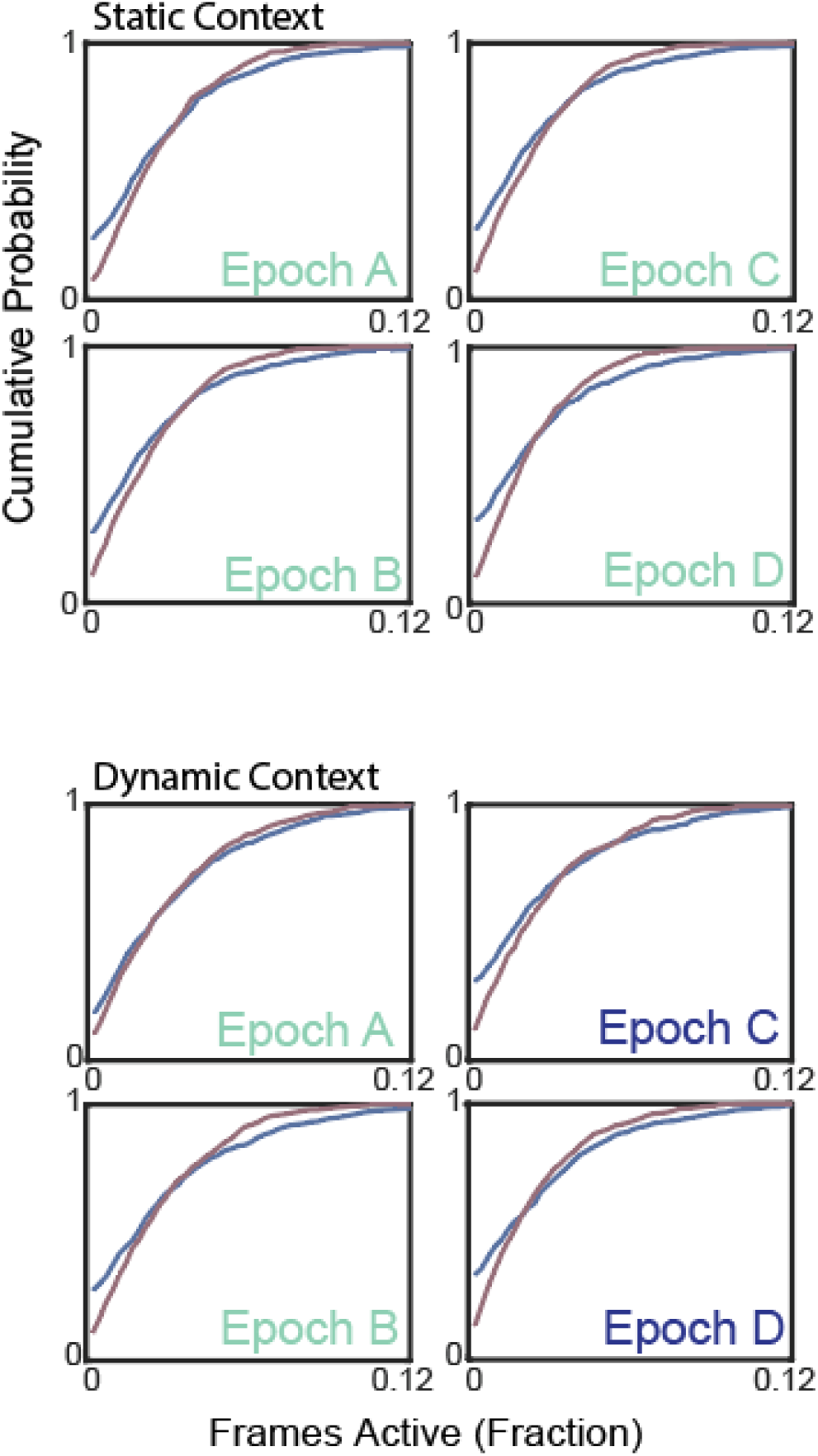
Dynamic changes in prefrontal network activity driven by social interaction regardless of context. **Top.** Cumulative probability function depicting the proportion of frames that each neuron was active during social (blue) and non-social frames (red) over the entire experiment (left) or during individual epochs (right). Social interaction was associated with an increase in both the proportion of highly active and inactive neurons in the static context experiment (Epoch A mean social 2.8+/-0.3% vs non-social 2.7+/-0.3%, p < 0.0001, KS Test; Epoch B mean social 2.6+/-0.2% vs non-social 2.6+/- 0.3%, p < 0.0001, KS Test; Epoch C mean social 2.3+/-0.2% vs non-social 2.3+/-0.3%, p < 0.0001, KS Test; Epoch D mean social 2.2+/-0.2% vs non-social 2.1+/-0.2% of frames, p < 0.0001, KS Test; n = 688 neurons from 8 mice. All comparisons nonsignificant by paired t-test; All comparisons p < 0.0001; KS test). **Bottom.** Cumulative distributions of activity for dynamic context experiment (Epoch A mean social 2.8+/-0.4% vs exploration 2.6+/-0.5%; NS by paired t test, p < 0.001, KS test; Epoch B mean social 2.3+/-0.4% vs non- social 2.1+/- 0.5%; NS by paired t test, p < 0.0001, KS test; Epoch C mean social 2.6+/-0.5% vs non-social 2.4+/-0.5%; NS by paired t test, p < 0.0001, KS test; Epoch D mean social 2.4+/-0.5% vs non-social 2.0+/- 0.4% of frames; n = 632 neurons from 8 mice).

**Supplementary Figure 2:**
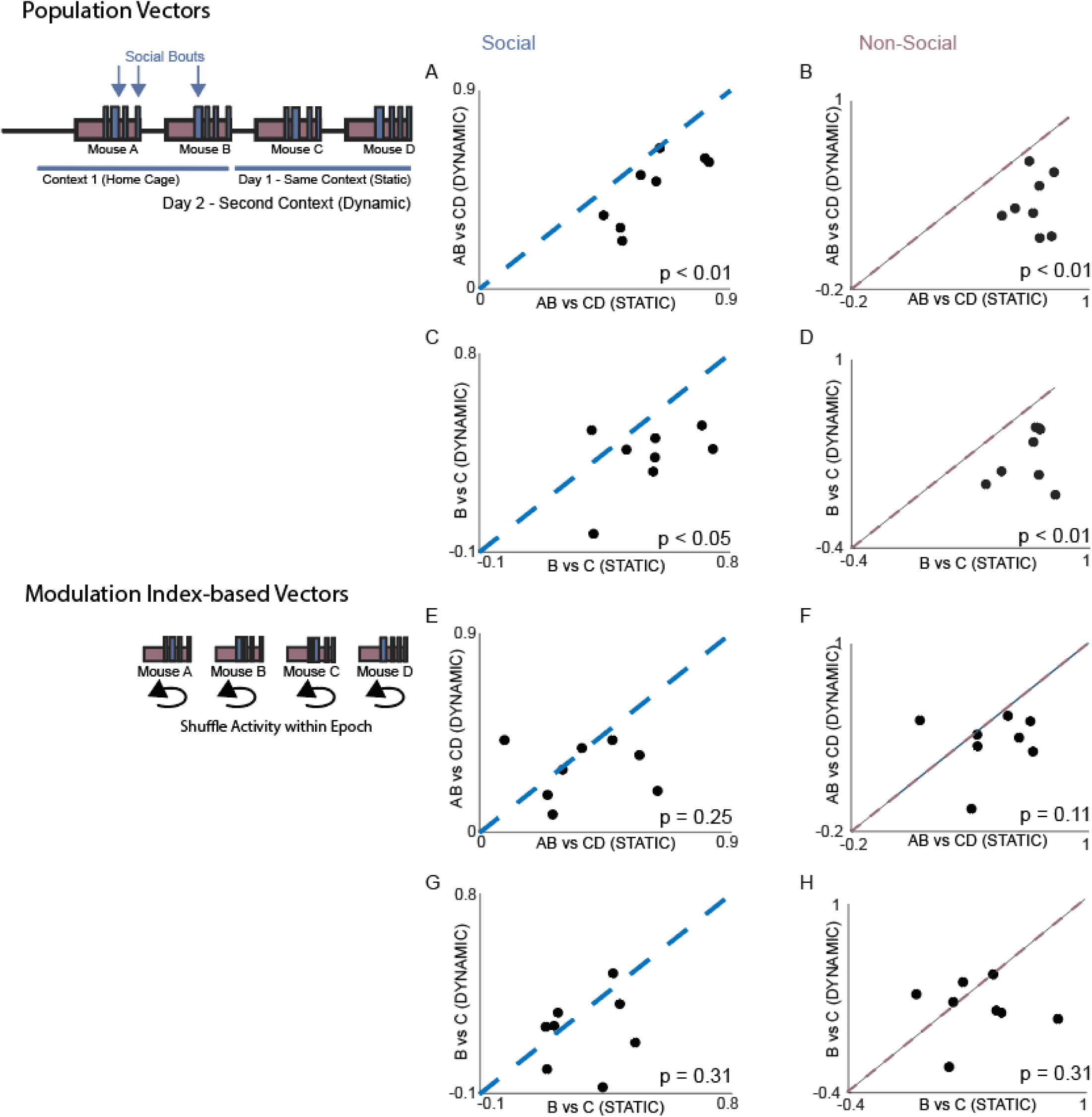
Socially modulated neurons are persistent despite context dependent remapping of prefrontal activity. We first calculated population vectors corresponding to the mean activity within each behavior by epoch. **A.** Scatterplot of the similarity between population vectors generated by averaging activity across social epochs A and B vs population vectors generated by averaging activity across social epochs C and D in the static context (x axis) and dynamic context (y-axis). Mean correlation static context 0.62+/-0.05 vs dynamic context 0.46+/-0.06, p < 0.01, signed-rank; n = 8 mice. **B.** Scatterplot of the similarity between population vectors generated by averaging activity across non-social epochs A and B vs population vectors generated by averaging activity across non-social epochs C and D in the static context (x axis) and dynamic context (y-axis). Mean correlation static context 0.72+/-0.03 vs dynamic context 0.35+/-0.06, p < 0.01, signed-rank; n = 8 mice. **C.** Scatterplot of the similarity between population vectors generated by averaging activity within social epochs B vs population vectors generated by averaging activity within social epoch C in the static context (x axis) and dynamic context (y-axis). Mean correlation static context 0.51+/-0.06 vs dynamic context 0.33+/-0.06, p < 0.05, signed-rank; n = 8 mice. **D.** Scatterplot of the similarity between population vectors generated by averaging activity within non-social epochs B vs population vectors generated by averaging activity within non-social epochs C in the static context (x axis) and dynamic context (y-axis). Mean correlation static context 0.64+/-0.05 vs dynamic context 0.28+/- 0.07, p < 0.01, signed-rank; n = 8 mice. We next calculated population vectors generated by calculating the Modulation Index for each neuron across each behavior and epoch. **E.** Scatterplot of the similarity between population vectors generated from the social modulation index generated from epochs A and B vs population vectors corresponding to the social modulation index generated from epochs C and D in the static context (x axis) and dynamic context (y-axis). Mean correlation static context 0.37+/-0.06 vs dynamic context 0.27+/-0.05, p = 0.25, signed-rank; n = 8 mice. **F.** Scatterplot of the similarity between population vectors generated from the non-social modulation index generated from epochs A and B vs population vectors corresponding to the non-social modulation index generated from epochs C and D in the static context (x axis) and dynamic context (y-axis). Mean correlation static context 0.37+/-0.07 vs dynamic context 0.22+/-0.07, p = 0.11, signed-rank; n = 8 mice. **G.** Scatterplot of the similarity between population vectors generated from the social modulation index generated from epoch B vs population vectors corresponding to the social modulation index generated from epoch C in the static context (x axis) and dynamic context (y-axis). Mean correlation static context 0.27+/-0.05 vs dynamic context 0.18+/-0.05, p = 0.31, signed-rank; n = 8 mice. **H.** Scatterplot of the similarity between population vectors generated from the non-social modulation index generated from epoch B vs population vectors corresponding to the non-social modulation index generated from epoch C in the static context (x axis) and dynamic context (y-axis). Mean correlation coefficient static context 0.27+/- 0.08 vs dynamic context 0.08+/-0.07, p = 0.31, signed-rank, n = 8 mice.

**Supplementary Figure 3:**
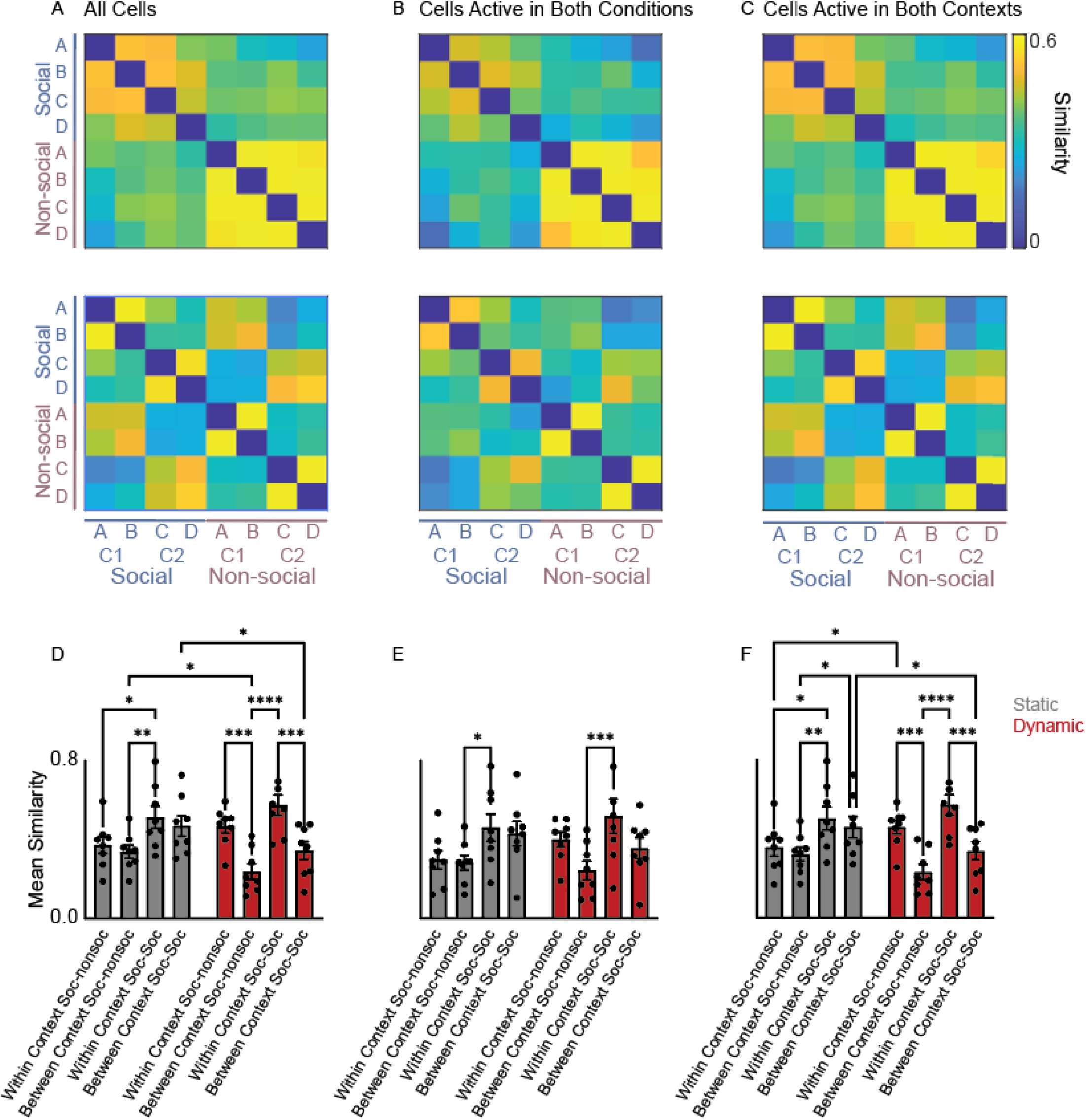
Context-dependent social encoding is not dependent on context-dependent activity. **A-C.** We examined the similarity of (correlation between) population activity vectors using either: all cells (A); only cells that were active within both vectors being compared (i.e., in both contexts) (B); only cells that were active for at least one behavioral condition (either social or nonsocial) in each context (context 1: epochs A & B; context 2: epochs C & D) (C). **D.** Bar graph of the similarities of (correlations between) population activity vectors for social or nonsocial behaviors in different contexts (gray and red bars correspond to the static and dynamic context experiments, respectively). In the dynamic context experiment, social vectors are more similar to social vectors from the same context than to cross-context social vectors; nonsocial vectors are also more similar to social vectors from the same context than to cross-context social vectors (2-way ANOVA, p = 0.69 for Experiment Day, p < 0.005 for for Comparison Type (social-social and social-nonsocial comparisons within or between context) and p < 0.01 for Experiment Day x Comparison Type, posthoc testing performed using Tukey’s multiple comparison test). **E.** Analogous to D, but for each comparison between a pair of population activity vectors, we excluded cells that were not active in both vectors (as in panel B). Though there remains a trend toward social vectors being more similar to within-context social vectors (compared to between-context comparisons) and non-social vectors also being more similar to within-context social vectors (compared to between-context comparisons) in the dynamic context experiment this is no longer significant (2-way ANOVA p = 0.88 for Experiment Day, p < 0.005 for Comparison Type (social-social and social-nonsocial comparisons within or between context) and p = 0.16 for Experiment Day x Comparison Type, posthoc testing performed using Tukey’s multiple comparison test, n = 6 mice). **F.** Analogous to D, but in this case we excluded cells that were not active in both contexts (similar to panel C; unlike E, cells were not necessarily active in each vector being compared). In the dynamic context experiment, social vectors are more similar to social vectors from the same context than the other context, and non-social vectors are also more similar to social vectors from the same context than the other context (2-way ANOVA p = 0.81 for Experiment Day (Static context vs Dynamic context), p < 0.005 for for Comparison Type (social-social and social-nonsocial comparisons within or between context)and p < 0.01 for Experiment Day x Comparison Type, posthoc testing performed using Tukey’s multiple comparison test).

**Supplementary Figure 4:**
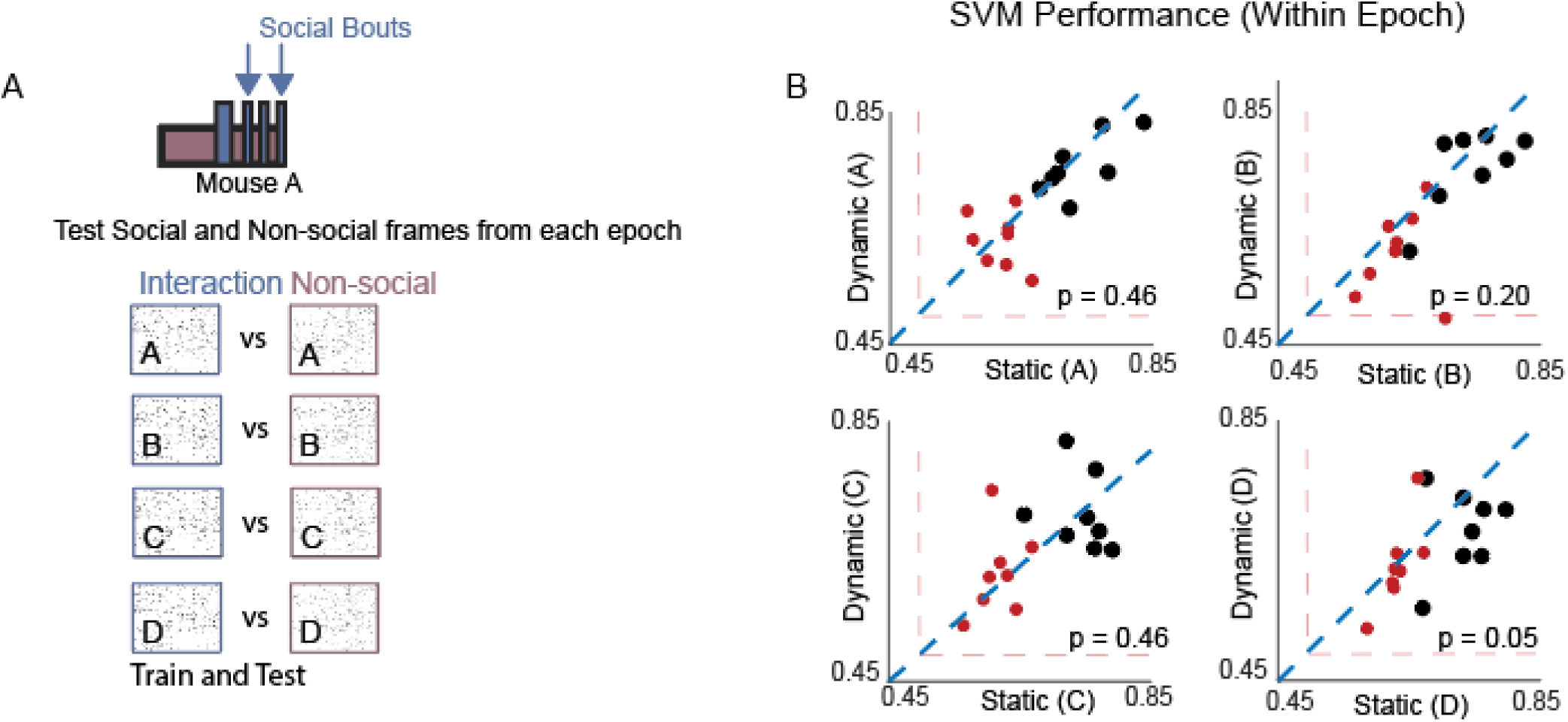
Linear Decoder discriminates social interaction regardless of context. **A.** A Support Vector Machine classifier was trained using binarized calcium data to discriminate frames corresponding to social interaction from non-social bouts within each epoch. **B.** Bargraph of the within-Epoch classifier performance for Epoch A-D. (Epoch A mean performance static context 73.7+/-1.8% vs dynamic context 71.9+/-1.6%, p = 0.46 signed-rank (shuffled values in red: 61.7+/- 1.1% static vs 60.7+/-1.5% dynamic); Epoch B mean performance static context 74.2+/-2.1% vs dynamic context71.9+/-2.2%, p = 0.20, signed-rank (shuffled values in red: 63.4+/-1.6% static vs 59.1% dynamic); Epoch C mean performance static context 74.9+/-1.4% vs dynamic context 73.0+/-2.4%, p = 0.46 signed-rank (shuffled values in red: 61.7+/-1.1% static vs 63.7+/-2.6% dynamic); Epoch D mean performance static context 73.6+/-1.4% vs dynamic context 67.5+/-2.2%, p = 0.05, signed-rank (shuffled values in red: 63.3+/-0.9% static vs 62.8+/-2.3% dynamic); n = 8 mice for all comparisons).

**Supplementary Figure 5:**
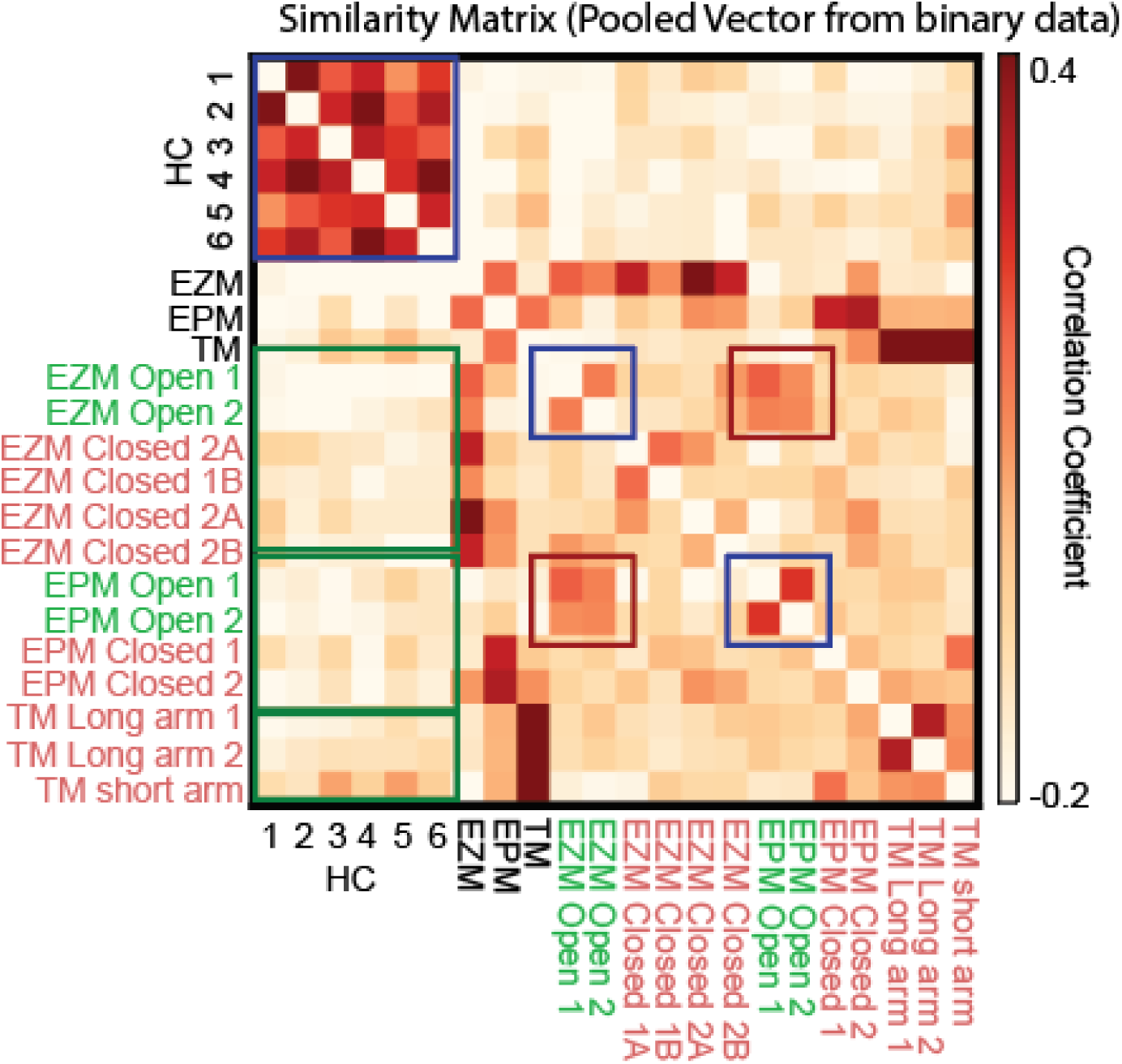
Context invariant representation of anxiety-related information. We generated a similarity matrix by computing correlations between population activity vectors corresponding to different epochs / specific arms. In this case population activity vectors consisted of all neurons pooled across mice. (HC vs HC epochs correlation coefficient 0.27, HC epochs vs all other epochs correlation coefficient -0.15, HC vs EZM correlation coefficient -0.16, HC vs EPM correlation coefficient -0.14, HC vs T-maze -0.09, all EZM subregions vs all EZM subregions correlation coefficient -0.01, all EPM subregions vs all EPM subregions correlation coefficient = 0.01, all T-maze subregions vs all T-maze subregions correlation coefficient 0.12, EZM open vs EPM open correlation coefficient 0.13, EZM Open vs T-Maze arms -0.08, EPM Open vs T-maze arms correlation coefficient -0.06, n = 633 neurons from 6 mice).

**Supplementary Figure 6:**
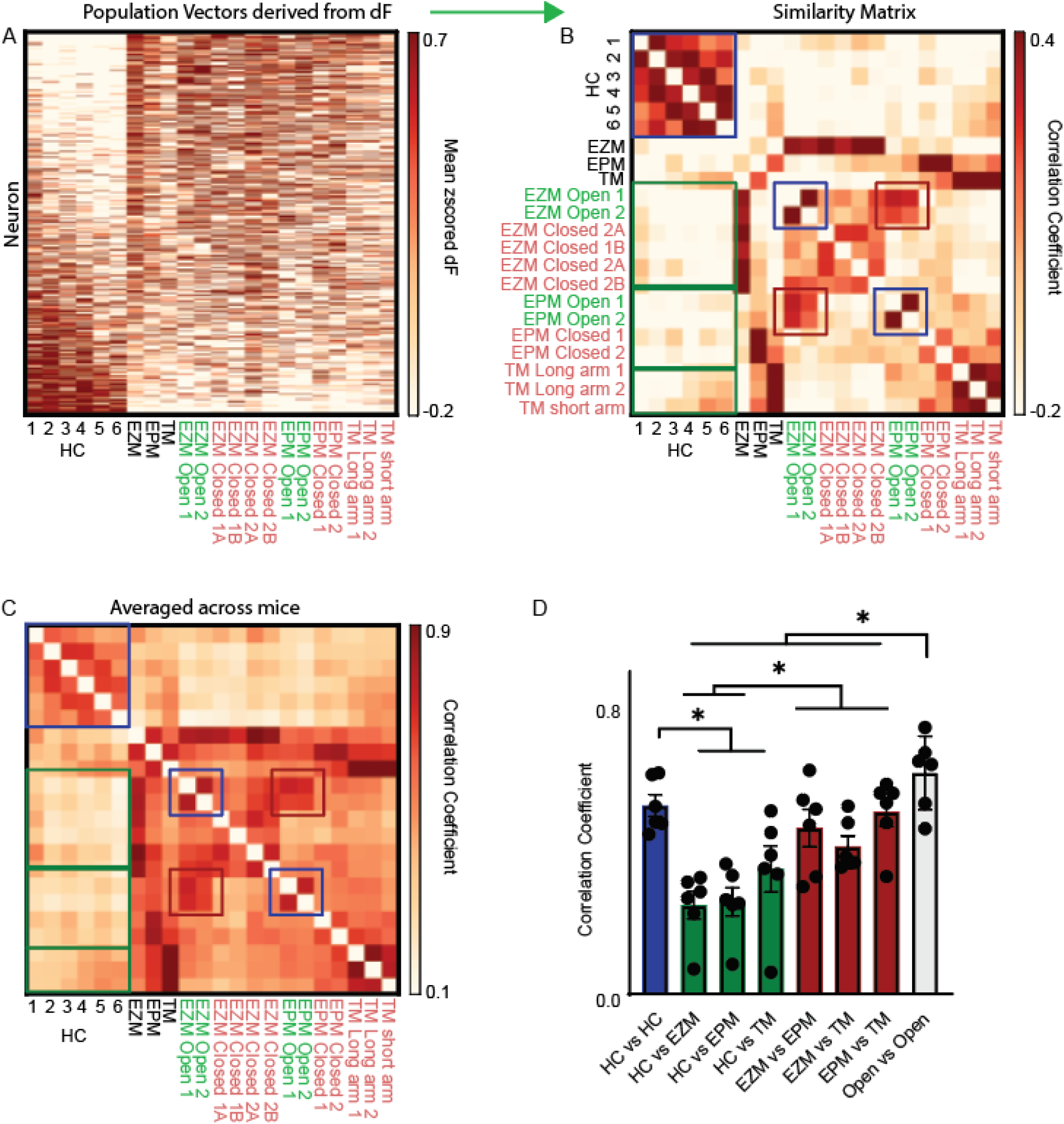
Persistence of ensemble representations of anxiety-related information across contexts. **A.** Behavior was subdivided into each home cage epoch and the time mice spent within each arm of the EZM, EPM, and T-Maze. EZM closed arms were further divided by the direction of travel (clockwise, CW; counter- clockwise, CCW). We generated population activity vectors for each epoch and arm by calculating the mean of the z-scored calcium trace for each neuron within each maze and each subregion (arm). Neurons were concatenated across all mice and then sorted the activity of neurons based on their average activity during home cage epochs; As in Figure 3 and 4 this revealed obvious differences between mPFC activity during home cage exploration and time spent in EZM, EPM, or T-Maze. **B.** We generated a similarity matrix by computing correlations between population activity vectors generated from averaged calcium traces corresponding to different epochs / specific arms. Population activity vectors were generated by concatenating all neurons from 6 mice (HC vs HC epochs correlation coefficient 0.31, HC epochs vs all other epochs correlation coefficient -0.18, HC vs EZM correlation coefficient -0.20, HC vs EPM correlation coefficient -0.18, HC vs T-maze -0.13, all EZM subregions vs all EZM subregions correlation coefficient 0.15, all EPM subregions vs all EPM subregions correlation coefficient = 0.06, all T-maze subregions vs all T-maze subregions correlation coefficient 0.37, EZM open vs EPM open correlation coefficient 0.24, EZM Open vs T-Maze arms-.12, EPM Open vs T-maze arms correlation coefficient -0.09, n = 633 neurons from 6 mice). **C.** Same as B but here we generated an averaged similarity matrix by computing correlations between population activity vectors corresponding to different epochs / specific arms for each mouse, then averaging the resulting 6 similarity matrices.. **D.** Bar graph depicting the averaged similarity between the 6 home cage epochs for each mouse (mean correlation coefficient 0.58+/-0.03), each context and the adjacent two home cage epochs (EZM vs HC mean correlation coefficient 0.28+/-0.04; EPM vs HC mean correlation coefficient 0.29+/-0.04; TMaze vs HC 0.38+/- 0.06), and the population activity underlying exploration of each of the three contexts (EZM vs EPM mean correlation coefficient 0.52+/-0.06, EZM vs Tmaze 0.46+/-0.03, EPM vs TMaze 0.57+/-0.04). ANOVA (p < 0.0001) with posthoc false discovery rate control using Benjamini, Krieger and Yekutieli). Asterisks represent discoveries.

**Supplementary Figure 7:**
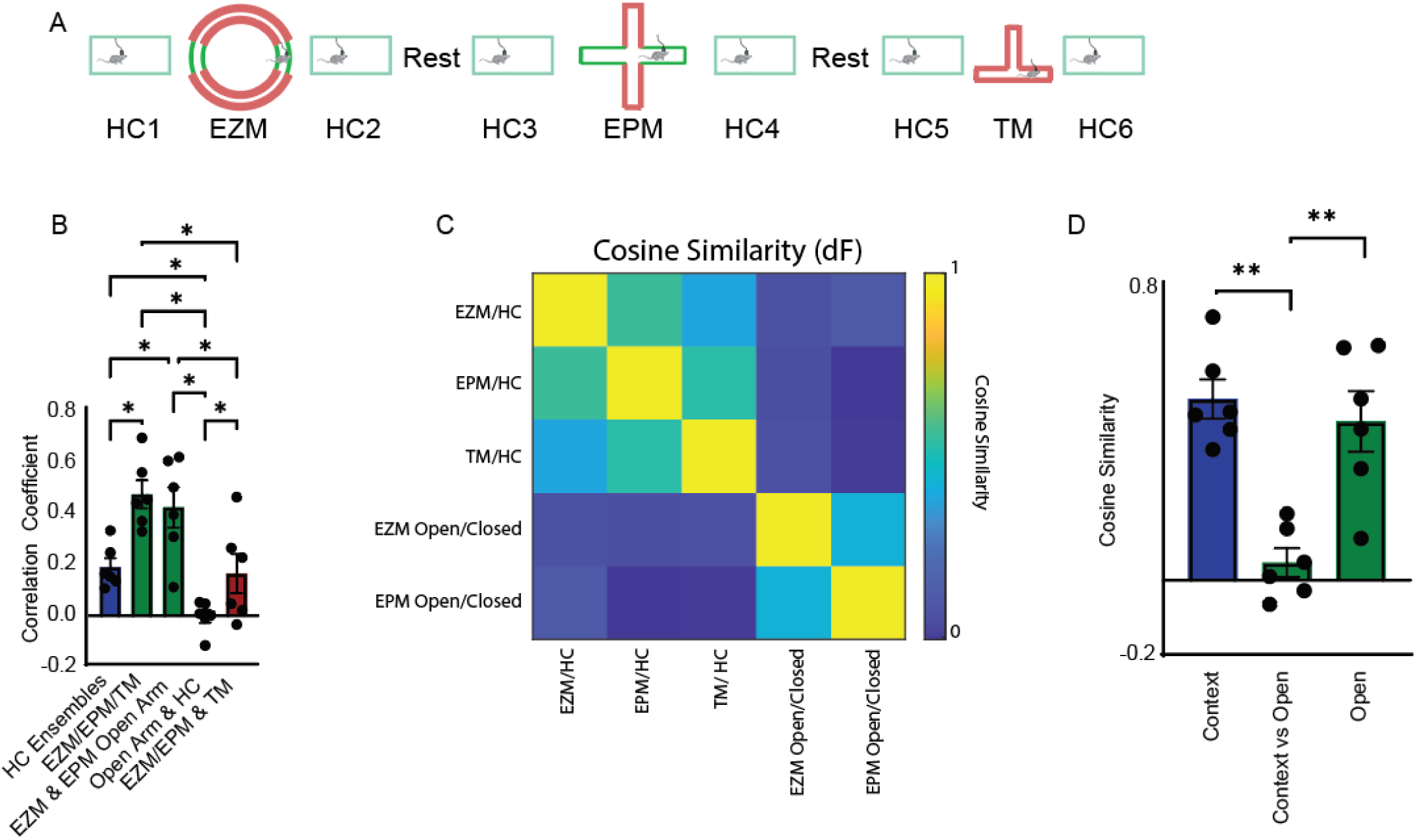
Orthogonal representations of anxiety- and context-related information. **A&B.** We compared modulation index derived ensembles underlying distinct behaviors and locations generated from the continuous calcium traces of each neuron. We generated HC vs HC ensembles by comparing activity in HC epoch 1vs2, 3vs4, and 5vs6. Comparison of the resulting vectors revealed little overlap (mean correlation coefficient 0.19+/-0.03). Context specific vectors were generated by comparing each context to adjacent periods of home cage exploration (mean correlation coefficient 0.48+/-0.06) and open arm specific ensembles for the EZM and EPM were generated by comparing activity during open arm exploration to closed arm exploration. Ensembles for open arm exploration were similar to ensembles generated for the open arm of the EPM (mean correlation 0.43+/-0.08). Open arm ensembles were distinct from HC vs HC ensembles (mean correlation coefficient -0.00+/-0.02). Finally, we quantified the similarity with which ensemble activity was modulated by exploration of the open arm in the EZM or EPM and the similarity with which ensemble activity was modulated as mice explored either of the two long arms of the Tmaze (mean correlation coefficient 0.17+/-0.07). Comparisons were made using ANOVA (p < 0.0001) with false discovery rate controlled using Benjamini, Krieger, and Yekutieli). Asterisks represent discovery. **C.** To determine the similarity of population activity underlying context and anxiety-related behavior, we calculated the cosine similarity between context-specific vectors and anxiety-related vectors. This data is plotted in a 5x5 matrix containing pairwise comparisons made between each of three context-specific vectors and each of the two anxiety-specific vectors. **D.** Bar graph depicting the mean cosine similarity within and between context- and anxiety-specific population vectors (mean cosine similarity context vs context: 0.49+/-0.05, mean cosine similarity context vs open arm: 0.05+/-0.04, mean cosine similarity open vs open: 0.43+/-0.08). RM ANOVA P < 0.001, posthoc Tukey’s multiple comparisons test.

**Supplementary Figure 8.**
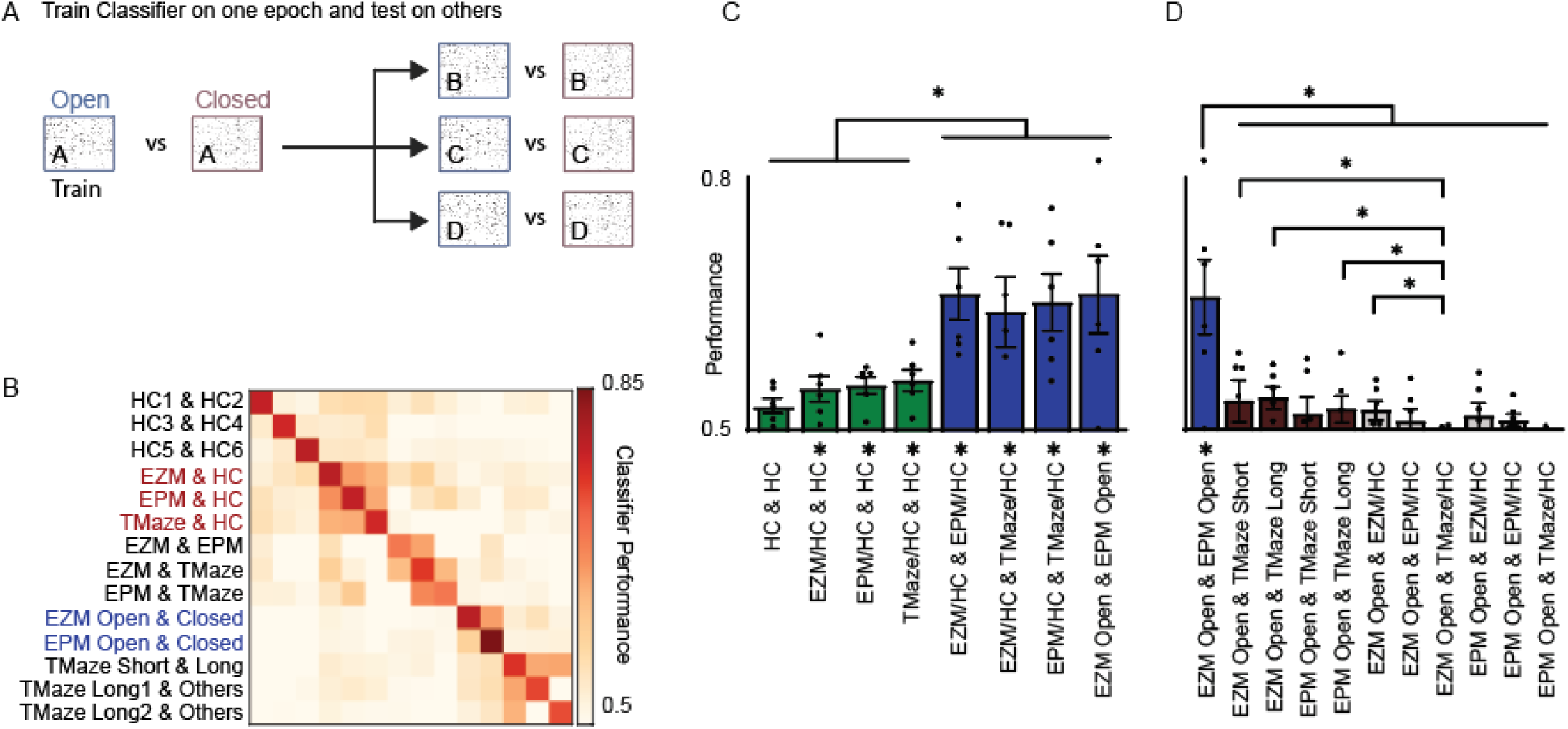
Decoding of behavioral states across context. **A.** We trained a linear classifier to distinguish frames corresponding to different mazes (ie home cage or EZM), epochs (home cage epoch 1 & 2), or subregions within specific mazes (ie EZM open & closed arms). In each case models were trained on binarized activity data corresponding to a given pair of behavioral regions or epochs, then tested the trained model on all 14 denoted comparisons (200 iterations, 0 hold out). Shuffled data was generated by circularly shuffling the ‘testing’ dataset, and reduced classifier performance to chance for all comparisons (not shown). **B.** 14x14 square matrix of classifier accuracy for all combinations of training and testing data. **C.** Bar graph of the mean performance of the SVM trained and tested on the corresponding pairs. Here ‘HC’ corresponds to pairs of home cage epochs adjacent to a single maze (ie ‘HC1 & HC2’), and X/HC indicates a model trained to distingish frames corresponding to exploration of ‘X’ from frames spent exploring the adjacent two home cage epochs. The classifier performed significantly above chance when trained to distinguish any of the three mazes from adjacent home cage epochs and then tested on any of the other mazes. Notably, the classifier was equally able to distinguish open arms from closed arms when trained on the EZM or EPM and tested on the opposing maze (HC & HC mean performance 0.53+/-0.01 (shuffled 0.50+/-0.01), EZM/HC & HC mean performance 0.54+/-0.02 (shuffled 0.50+/-0.01), EPM/HC & HC mean performance 0.55+/-0.01 (shuffled 0.50+/-0.01), TMaze/HC & HC mean performance 0.56+/-0.01 (shuffled 0.51+/-0.01), EZM/HC & EPM/HC mean performance 0.66+/-0.03 (shuffled 0.51+/-0.00), EZM/HC & TMaze/HC mean performance 0.64+/- 0.04(shuffled 0.52+/-0.01), EPM/HC & TMaze mean performance 0.65+/-0.03 (shuffled 0.49+/-0.02), EZM Open & EPM Open mean performance 0.66+/-0.05 (shuffled 0.51+/-0.01). Data from panels C and D are subsets of same analysis; p < 0.0001 for ensemble comparison, p =< 0.01 for shuffling, and p < 0.0001 for interaction; 2 WAY RM ANOVA with posthoc control of false discovery using two stage linear procedure of Benjamini, Krieger, and Yekutieli. Shuffled data not shown but * below x-axis indicate comparisons that differ significantly from shuffled data. **D.** Bar graph of within-context, and within-context vs context comparisons. The classifier performed above chance only when trained to distinguish the open arm of one maze and tested on the opposing maze. All other comparisons were indistinguishable from shuffled data. (EZM Open & TMaze Short mean performance 0.53+/-0.03 (shuffled 0.51+/-0.01), EZM Open & TMaze Long mean performance 0.54+/-0.01 (shuffled 0.50+/-0.01), EPM Open & TMaze Short mean performance 0.52+/-0.02 (shuffled 0.49+/-0.01), EZM Open & EZM/HC 0.52+/-0.01 (shuffled 0.51+/-0.01), EZM Open & EPM/HC mean performance 0.51+/-0.02 (shuffled 0.50+/- 0.01), EZM open & TMaze mean performance 0.48+/-0.01 (shuffled 0.51+/-0.01), EPM Open & EZM/HC mean performance 0.52+/-0.02 (shuffled 0.50+/-0.01), EPM Open & EPM/HC mean performance 0.51+/-0.01 (shuffled 0.50+/-0.01), EPM Open & TMaze/HC mean performance 0.48+/-0.01 (shuffled 0.49+/-0.01)). Shuffled data not shown but * below x-axis indicate comparisons that differ significantly from shuffled data.

## References

1. A. C. Brumback et al., Identifying specific prefrontal neurons that contribute to autism-associated abnormalities in physiology and social behavior. Mol Psychiatry 23, 2078–2089 (2018).

2. 2. A. Selimbeyoglu et al., in Science translational medicine. (American Association for the Advancement of Science, 2017), vol. 9, pp. eaah6733.

3. D. R. Levy et al., Dynamics of social representation in the mouse prefrontal cortex. Nature neuroscience, (2019).

4. A. T. Lee et al., VIP Interneurons Contribute to Avoidance Behavior by Regulating Information Flow across Hippocampal-Prefrontal Networks. Neuron 102, 1223–1234.e1224 (2019).

5. A. Adhikari, M. A. Topiwala, J. A. Gordon, Single units in the medial prefrontal cortex with anxiety-related firing patterns are preferentially influenced by ventral hippocampal activity. Neuron 71, 898–910 (2011).

6. S. Ciocchi, J. Passecker, H. Malagon-Vina, N. Mikus, T. Klausberger, Brain computation. Selective information routing by ventral hippocampal CA1 projection neurons. Science 348, 560–563 (2015).

7. N. Padilla-Coreano et al., Direct Ventral Hippocampal-Prefrontal Input Is Required for Anxiety-Related Neural Activity and Behavior. Neuron 89, 857–866 (2016).

8. M. Murugan et al., Combined Social and Spatial Coding in a Descending Projection from the Prefrontal Cortex. Cell 171, 1663–1677 e1616 (2017).

9. D. K. Lee et al., Reduced sociability and social agency encoding in adult Shank3-mutant mice are restored through gene re-expression in real time. Nature Neuroscience 24, 1243–1255 (2021).

10. B. Liang et al., Distinct and Dynamic ON and OFF Neural Ensembles in the Prefrontal Cortex Code Social Exploration. Neuron 100, 700–714.e709 (2018).

11. J. H. Jennings et al., Interacting neural ensembles in orbitofrontal cortex for social and feeding behaviour. Nature 565, 645–649 (2019).

12. N. A. Frost, A. Haggart, V. S. Sohal, Dynamic patterns of correlated activity in the prefrontal cortex encode information about social behavior. PLoS Biol 19, e3001235 (2021).

13. R. C. deCharms, M. M. Merzenich, Primary cortical representation of sounds by the coordination of action- potential timing. Nature 381, 610–613 (1996).

14. E. Vaadia et al., Dynamics of neuronal interactions in monkey cortex in relation to behavioural events. Nature 373, 515–518 (1995).

15. P. Xu et al., Pattern decorrelation in the mouse medial prefrontal cortex enables social preference and requires MeCP2. Nature communications 13, (2022).

16. O. Yizhar et al., Neocortical excitation/inhibition balance in information processing and social dysfunction. Nature 477, 171–178 (2011).

17. 17. D. Amodio, C. Frith, Meeting of minds: the medial frontal cortex and social cognition. Nature reviews. Neuroscience 7, (2006).

18. A. Adhikari, M. A. Topiwala, J. A. Gordon, Synchronized activity between the ventral hippocampus and the medial prefrontal cortex during anxiety. Neuron 65, 257–269 (2010).

19. M. Cunniff, E. Markenscoff-Papadimitriou, J. Ostrowski, J. Rubenstein, V. Sohal, Altered hippocampal-prefrontal communication during anxiety-related avoidance in mice deficient for the autism-associated gene Pogz. eLife 9, (2020).

20. E. Likhtik, J. M. Stujenske, M. A. Topiwala, A. Z. Harris, J. A. Gordon, Prefrontal entrainment of amygdala activity signals safety in learned fear and innate anxiety. Nat Neurosci 17, 106–113 (2014).

21. V. Hok, E. Save, P. P. Lenck-Santini, B. Poucet, Coding for spatial goals in the prelimbic/infralimbic area of the rat frontal cortex. Proc Natl Acad Sci U S A 102, 4602–4607 (2005).

22. J. Sauer, S. Folschweiller, M. Bartos, Topographically organized representation of space and context in the medial prefrontal cortex. Proceedings of the National Academy of Sciences of the United States of America 119, (2022).

23. J. M. Hyman, L. Ma, E. Balaguer-Ballester, D. Durstewitz, J. K. Seamans, Contextual encoding by ensembles of medial prefrontal cortex neurons. Proc Natl Acad Sci U S A 109, 5086–5091 (2012).

24. V. Jovanovic, A. Fishbein, L. de la Mothe, K. Lee, C. Miller, Behavioral context affects social signal representations within single primate prefrontal cortex neurons. Neuron 110, (2022).

25. E. A. Mukamel, A. Nimmerjahn, M. J. Schnitzer, Automated analysis of cellular signals from large-scale calcium imaging data. Neuron 63, 747–760 (2009).

26. Y. Sakurai, How do cell assemblies encode information in the brain? Neurosci Biobehav Rev 23, 785–796 (1999).

27. Y. Sakurai et al., Multiple Approaches to the Investigation of Cell Assembly in Memory Research-Present and Future. Front Syst Neurosci 12, 21 (2018).

28. G. Buzsaki, Neural syntax: cell assemblies, synapsembles, and readers. Neuron 68, 362–385 (2010).

29. L. Carrillo-Reid, W. Yang, Y. Bando, D. S. Peterka, R. Yuste, Imprinting and recalling cortical ensembles. Science 353, 691–694 (2016).

30. D. J. Cai et al., A shared neural ensemble links distinct contextual memories encoded close in time. Nature 534, 115–118 (2016).

31. Y. Ziv et al., Long-term dynamics of CA1 hippocampal place codes. Nat Neurosci 16, 264–266 (2013).

32. M. S. Fustiñana, T. Eichlisberger, T. Bouwmeester, Y. Bitterman, A. Lüthi, State-dependent encoding of exploratory behaviour in the amygdala. Nature 592, 267–271 (2021).

33. y. Li, Neuronal Representation of Social Information in the Medial Amygdala of Awake Behaving Mice: Cell. (2021).

34. R. Remedios et al., Social behaviour shapes hypothalamic neural ensemble representations of conspecific sex. Nature 550, 388–392 (2017).

35. K. Jantzen, O. Oullier, M. Marshall, F. Steinberg, J. Kelso, A parametric fMRI investigation of context effects in sensorimotor timing and coordination. Neuropsychologia 45, (2007).

36. J. Peters, I. Daum, E. Gizewski, M. Forsting, B. Suchan, Associations evoked during memory encoding recruit the context-network. Hippocampus 19, (2009).

37. E. Mankin, G. Diehl, F. Sparks, S. Leutgeb, J. Leutgeb, Hippocampal CA2 activity patterns change over time to a larger extent than between spatial contexts. Neuron 85, (2015).

38. M. W, H. ME, C. DJ, The brain in motion: How ensemble fluidity drives memory-updating and flexibility. eLife 9, (2020).

39. E. Miller, J. Cohen, An integrative theory of prefrontal cortex function. Annual review of neuroscience 24, (2001).

40. B. Barak, G. Feng, Neurobiology of social behavior abnormalities in autism and Williams syndrome. Nat Neurosci 19, 647–655 (2016).

41. M. L. Phillips, H. A. Robinson, L. Pozzo-Miller, Ventral hippocampal projections to the medial prefrontal cortex regulate social memory. eLife 8, e44182 (2019).

42. V. Torres-Lista, L. Giménez-Llort, Vibrating Tail, Digging, Body/Face Interaction, and Lack of Barbering: Sex- Dependent Behavioral Signatures of Social Dysfunction in 3xTg-AD Mice as Compared to Mice with Normal Aging. Journal of Alzheimer’s disease : JAD 69, (2019).

43. A. Arrant, A. Filiano, B. Warmus, A. Hall, E. Roberson, Progranulin haploinsufficiency causes biphasic social dominance abnormalities in the tube test. Genes, brain, and behavior 15, (2016).

44. N. Ghoshal, J. Dearborn, D. Wozniak, N. Cairns, Core features of frontotemporal dementia recapitulated in progranulin knockout mice. Neurobiology of disease 45, (2012).

45. T. Chow, M. Mendez, Goals in symptomatic pharmacologic management of frontotemporal lobar degeneration. American journal of Alzheimer’s disease and other dementias 17, (2002).

